# Food odours associated with conspecific corpses cause food avoidance in an invasive ant

**DOI:** 10.1101/2023.05.16.540952

**Authors:** Thomas Wagner, Tomer J. Czaczkes

**Affiliations:** Animal Comparative Economics Laboratory, Faculty of Biology and Preclinical Medicine, University of Regensburg, Universitaetsstrasse 31,93053 Regensburg, Germany

**Keywords:** Bait avoidance, learning, negative stimulus, odour association, corpse avoidance

## Abstract

Invasive ants, such as *Linepithema humile* (the Argentine ant), pose a global threat, necessitating a better understanding of their behaviour in order to improve management strategies. Traditional eradication methods, including baiting, have had limited success, but the causes of control failure are not always clear. Here we propose that ants may learn to avoid toxic baits in part due to their association with ant corpses. Ants were tested on a Y-maze after exposure to scented corpses or dummies. 69% (n = 64) of ants avoided branches bearing the scent of scented corpses. At a collective level, colonies neglected food sources associated with scented ant corpses in favour of a food source with a novel odour, with only 42% (n = 273) of foragers feeding from the corpse-scent associated food source. However, if corpses were produced by feeding ants scented toxicant, focal ants encountering these corpses did not avoid the corpse-associated scent on a Y maze (53%, n = 65). Moreover, in a dual-feeder test, ants did not avoid feeding at food sources scented with an odour associated with conspecific corpses. The study demonstrates that conspecific corpses can act as a negative stimulus for *Linepithema humile*, leading to avoidance of odours associated with corpses, which can lead to potential avoidance of toxic baits. Why the more realistic Y-maze trial with corpses of ants that had ingested the toxicant elicited no avoidance is unclear: it may be due to weaker odour cues from ingested food, or a counterbalancing of the negative corpse stimulus by the positive presence of food remains on the corpse. Nonetheless, this study demonstrates that conspecific corpses act as a negative stimulus for ants, and this should be kept in mind when planning control efforts. A simple solution to this issue would be adding odours to baits, and cycling baits between treatments. This would disrupt the association between bait odour and corpses.

## Introduction

Invasive ant species can have a significant impact on ecosystems and economies alike, making it crucial to understand their behaviour and find effective management strategies to reduce their consequences (1–3). *Linepithema humile* (Mayr, 1868), commonly known as the Argentine ant, is a highly destructive ant species that causes significant ecological and economic damage globally (4), and is the most widely distributed invasive ant species in Europe (5). Invasion by *L. humile* leads to a significant reduction in invertebrate diversity, and can also have an impact on vertebrates (6–8). In addition, *L. humile* can be an important agriculture pest by enhancing Hemipteran populations, which then increases the likelihood of fungal and viral infections (9).

Effective management of invasive ants can have large beneficial environmental outcomes, especially in terms of restoring arthropod biodiversity (10). Unfortunately, invasive ants are difficult to control: two-thirds of *L. humile* eradication attempts have failed (10). The current gold standard approach for eradication or control of ants is to use baits with a toxicant that can be shared with the entire colony by the foragers before the effects are realised, allowing toxicants to reach the brood and queen(s) (10). Even so, the success rate of eradication attempts is low (10,11).

We propose that one reason for control failure may be that ants learn to avoid bait-associated cues because they are also associated with ant corpses, created during control efforts. This hypothesis assumes that nestmate corpses act as an *unconditional negative stimulus* (12,13). It has been shown that the scent of food that ants have consumed can be detected by nestmates that encounter them, even without active food sharing (14–17). The scent of a toxicant-laced bait, remaining on ants after they have died, may lead to an aversion to the bait among the living ants that encounter these corpses, causing them to avoid the scent or taste of the bait. Ant corpses are a potential threat to the survival of colonies due to the possibility of infection (18). To protect a colony from pathogens, it is crucial for ants to distinguish between living and dead ants. Recognition of corpses is likely based on olfactory cues (19). In *L. humile*, the presence of iridomyrmecin and dolichodial (ant-produced compounds that adhere to the cuticle) on live ants disappears within 40 minutes of death, which then triggers necrophoresis behaviour from nestmates (20). Studies have demonstrated that ant corpses can inhibit foraging or nesting. For example, it was shown that the presence of nestmate corpses can cause ants to reject otherwise suitable nesting sites (21,22) and refuse dumps can deter ants from foraging (23,24).

It is thus reasonable to expect that conspecific corpses may represent an *unconditional negative stimulus*. *L. humile* has a formidable olfactory memory and is able to rapidly form long-lasting associations between odours or flavours and a food reward (25–27). Nevertheless, it is not clear how well *L. humile* learns negative associations. Wagner et al. (26) showed that using quinine as a negative stimulus does not reinforce learning when a positive stimulus (sucrose reward) was already presented. However, *L. humile* preferred to choose “low-risk” paths to food sources after they discovered a competitor species or formic acid on a “high risk” path (28), and other ant species rapidly form negative olfactory associations, and avoid odours associated with, for example, a bitter quinine solution (29).

To the best of our knowledge, no previous studies have investigated whether exposure to conspecific corpses affects the feeding preferences and decision-making of foraging ants. We hypothesize that *L. humile* workers can form a negative association between the odour of a food source and the presence of conspecific corpses which fed on the same food source, or died in the food sourcés vicinity, and hence have the food scent on their bodies. Specifically, we investigated the impact of exposure to scented nestmate corpses on foraging behavior, and to determine whether this exposure leads to avoidance of high-quality food sources or areas.

## Materials and Methods

### Colony maintenance

*Linepithema humile* ants were collected in 2021 from Girona, Spain and were all part of the same main European supercolony (30). Colony fragments (henceforth colonies), consisting of one or more queens and 300-800 workers, were kept in plastic foraging boxes (32.5 x 22.2 x 11.4cm) with plaster of Paris on the bottom. No aggression between ants from different colony fragments was observed. The walls were coated in fluon to prevent escape. Each box contained several 15mL plastic tubes, covered in transparent red foil, partly filled with water and plugged with cotton, for use as nests. The ants were maintained on a 12:12 light:dark cycle at room temperature (21-25 °C) and provided with water *ad libitum*. Colonies were fed for three days with *ad libitum* 0.5M sucrose solution and freeze-killed *Drosophila melanogaster*, and deprived of food for four days prior to testing. All experiments were conducted between October 2021 and June 2022.

### Solutions and odours

1M sucrose solutions (Südzucker AG, Mannheim, Germany), were used as a reward during training for all experiments. Scented paper overlays, used during experiments to provide environmental odours, were stored for at least 1 week prior to the experiments in airtight plastic boxes (19.4 x 13.8 x 6.6cm) containing a glass petri dish with 500µL of either strawberry or apple food flavouring, or blackcurrent and orange food flavouring (Seeger, Springe, Germany). For experiments where flavoured food was used, 1µL of the respective flavouring was added per 1mL of 1M sucrose solution.

### Corpses and dummies

One day prior to the start of experiments, 24 ants were frozen for 60 min in a -20°C freezer. These ant corpses were scented by placing them at air temperature for 24 hours into an airtight petri dish (80x15mm, sealed with Parafilm©) containing a small metal plate with 200µL of strawberry or apple food flavouring, or with blackcurrent or orange flavouring. *L. humile* can identify ants as being dead within 40 minutes of death (20), so 24 hours was considered more than sufficient for corpses to be identified as such by nestmates. Note that we did not attempt to test ants only with corpses from their own fragments. However, all colony fragments were sourced from the same location at the same time, and showed no aggression between each other.

24 dummies were created by cutting black electrical wire (mm diameter and 2.0-2.5mm length) to visually simulate corpses, and provide a similar obstacle to movement. Dummies were scented identically to the corpses. We chose to use plastic-coated wire, rather than glass beads, as we considered plastic to be more similar to the ants’ surface in terms of how readily it would absorb odours. Note that the corpses and the dummies had no physical contact with the food flavourings, but were exposed to the scented air in the sealed petri dish.

### Experiment 1 – Testing the effect of corpse-scent association on odour preference using scented corpses

Here we investigate whether ants are capable of forming an association between an odour and dead nestmates (corpses) in a Y-maze test setting. We predicted that ants exposed to scented corpses would avoid the smell associated with the corpses, but ants exposed to scented dummies would not.

12 corpses (treatment) or dummies (control) were placed onto a round plastic arena surrounded by a water moat (arena: 7cm diameter, inner platform: 4cm diameter, see figure 1A) using clean, soft forceps. A colony was connected via a drawbridge to the round arena and a single ant was allowed to enter the platform in the arena via the drawbridge. The drawbridge was then removed, trapping the ant in the arena together with the scented corpses or scented dummies (e.g. orange). After 90 seconds, or at least 10 contacts between the ant and the corpses or dummies, a second drawbridge was added to let the ant leave the arena and enter a Y-maze (see fig. 1B).

**Figure 1:**
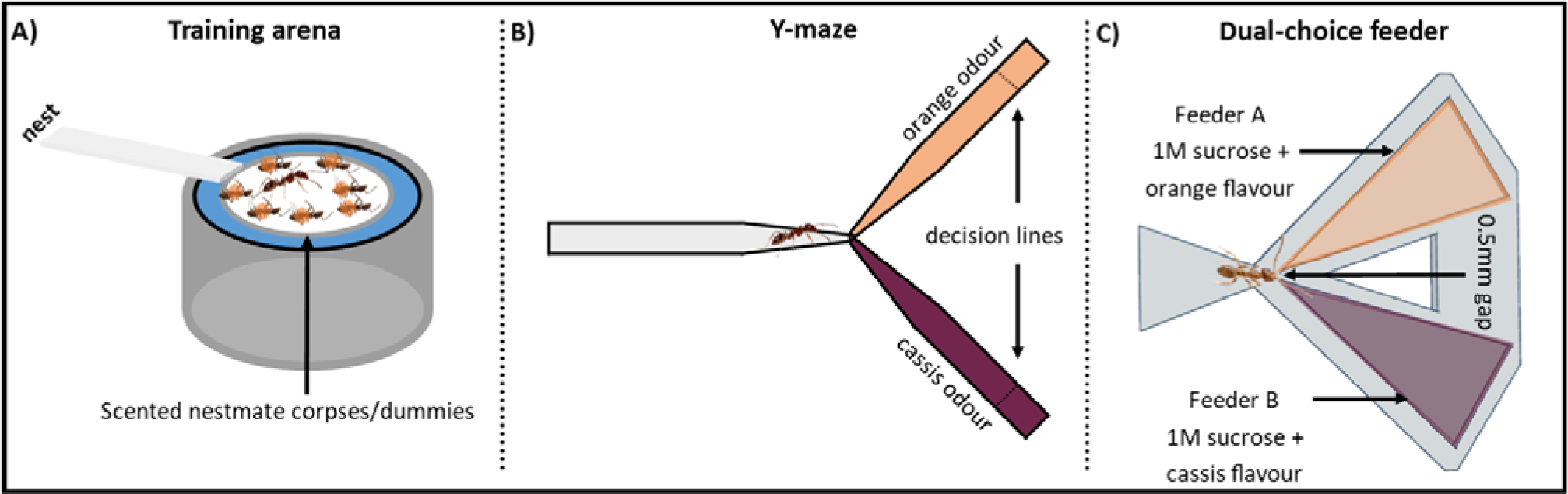
Setup used for experiments 1-3: A) Ants were confined to an encounter arena, containing scented dummies or nestmate corpses (experiments 1 & 3), or corpses of ants that had fed on scented toxicant (experiment 2). After 90s or 10 contacts, in experiments 1 & 2, the ants’ odour preference was then tested on a Y-maze with scented paper overlays, the colour referring to the two different odours. We noted which decision line was crossed. In experiment 3 ants were tested on a dual-choice feeder (C): Ants enter via a constriction, encountering two adjacent wells, containing differently-flavoured sucrose solution. The ants first choice, time to first feeding interruption, and total feeding time from each well was tested. Note that the two well tips are so close as to ensure the ant contacts both wells simultaneously (ant image to scale).

The stem of the Y-maze (i.e. the part through which the ant enters the Y-maze) was covered in an unscented paper overlay, one arm (e.g. left past the bifurcation point) was covered in a scented overlay with the scent matching the scent of the corpses/dummies (e.g. orange), and the other arm was covered in a neutral-scented overlay (novel odour e.g. blackcurrent). We recorded the ant’s final decision (crossing a line 2cm from the arm end, see fig. 1B). The position of the odour associated with the corpses or dummies was systematically varied between ants. The Y-maze was cleaned with ethanol after each ant. 16 ants were tested sequentially using the same ant corpses, followed by 16 tested with dummies of the same smell. The next day, the same procedure was performed with the opposite odour (e.g. blackcurrent). The negative-associated odour, and the sequence of corpses/dummies were switched alternately. 32 ants were tested per day, testing in total 128 ants from 4 colonies.

### Experiment 2 – Testing the effect of corpse-scent association on odour preference using corpses killed with a scented toxicant

Experiment 1 found that the ants did indeed avoid the corpse-associated odour, but not the dummy associated odour. The aim of this experiment was to test whether the same applied when corpses were infused with scent from just feeding on a flavoured toxic food source. Two days prior to start of the experiment hydrogel beads (31,32) were placed into a solution containing 7.6µl Spintor^®^ (containing 44% Spinosad) and 200µl food flavour (either blackcurrent or orange) in 100ml 1M sucrose. One day prior the start of the experiment, 24 ants were fed on these hydrogel beads in a petri dish. After 8h all ants were dead, and the corpses were placed in a clean petri dish. The next day (∼16h later) these ants were used as corpses in the behavioural trials. The procedure of the training and testing was identical to experiment 1. Note that a control treatment with dummy ants was not possible in this experiment, as consumption of the flavoured toxicant is required for odour presentation during the experiment. 16 ants were tested per day, testing in total 64 ants from 4 colonies over 4 days on corpses.

### Experiment 3 – Testing the effect of corpse-associated odour on feeding preference

Here we asked whether ants respond differently to food flavoured to match the scented nestmates corpses or scented dummies. Furthermore, we tested if a lower starvation level (2 days) changed the response of the ants compared to the standard 4 days. The idea here was that extremely hungry ants may be too motivated to feed at the first food encountered to show preference between the two options.

Ants encountered corpses or dummies, scented by storing with food flavourings for 24 hours, in the round plastic arena as described in experiment 1. After this training the ant was allowed to enter a drawbridge leading to a ‘dual-choice’ feeder, made of 3D printed resin (see fig. 1C). This feeder consisted of two triangular wells, each filled with a different solution (scented 1M sucrose). The tips of the wells were 0.5mm apart, smaller than the length of one *L. humile* antenna. This ensured that, regardless of which solution is contacted first, the ant will almost immediately also come into contact with the alternative solution, enabling an informed choice between the two solutions (see figure 1C). We noted the time spent feeding from each feeder.

The feeder side containing the corpse-associated flavoured food was systematically varied between ants. 32 ants were tested per day, testing in total 127 ants from 4 colonies. 64 ants were trained on dummies and 63 on corpses. Tested ants were permanently excluded from the colony, to prevent scented food being returned to their nestmates.

### Experiment 4 – Can the presence of scented corpses drive collective food choice?

The purpose of this experiment was to raise the results of the individual level experiments to the collective level, quantifying collective foraging at a high resolution over the first hour of feeding, and at a lower resolution over 24 hours. Data were split into the first 60min after food presentation, and the following 23h. This is because in a laboratory setting, satiation at *ad libitum* feeders often occurs within the first hour (33).

Hydrogel beads were prepared as in experiment 2, but without the toxicant. A round platform (9cm diameter, 0.2cm thick), raised on 4 plastic pillars (0.3cm diameter, 5.5m height), connected to a raised a plastic bridge, was placed into the colony box. The pillars were coated with fluon to ensure that the only access to the platform was via the bridge. Two hydrogel beads, flavoured to match the scent of the corpses, were placed into a 0.1mm deep trough on one side of the round platform (see fig. 2), and two other hydrogel beads with the other flavour were placed on the other side. These beads were cut in half to prevent them rolling (32). 12 scented corpses, prepared as in experiment 1, were distributed over the bridge (see fig. 2). Ants were then allowed free access to the feeders via the bridge. The narrow bridge ensured that all ants encountered most of the corpses. Ants were allowed to recruit and feed on the hydrogel breads for 24h. Images were recorded automatically by an infrared-sensitive camera with infrared flash (Pi NoIR module 2) connected to a microcomputer (Raspberry Pi 4). An image was taken every 5 minutes for the first 60 minutes of the experiment (corresponding to the major feeding and recruitment stage) and thereafter every hour for 24h. From the images, we counted the number of ants within 5mm of each feeder. The feeder side and the flavour containing the negative associated hydrogel beads was systematically varied between colonies. We tested 16 colonies, and each colony was only tested once.

**Figure 2:**
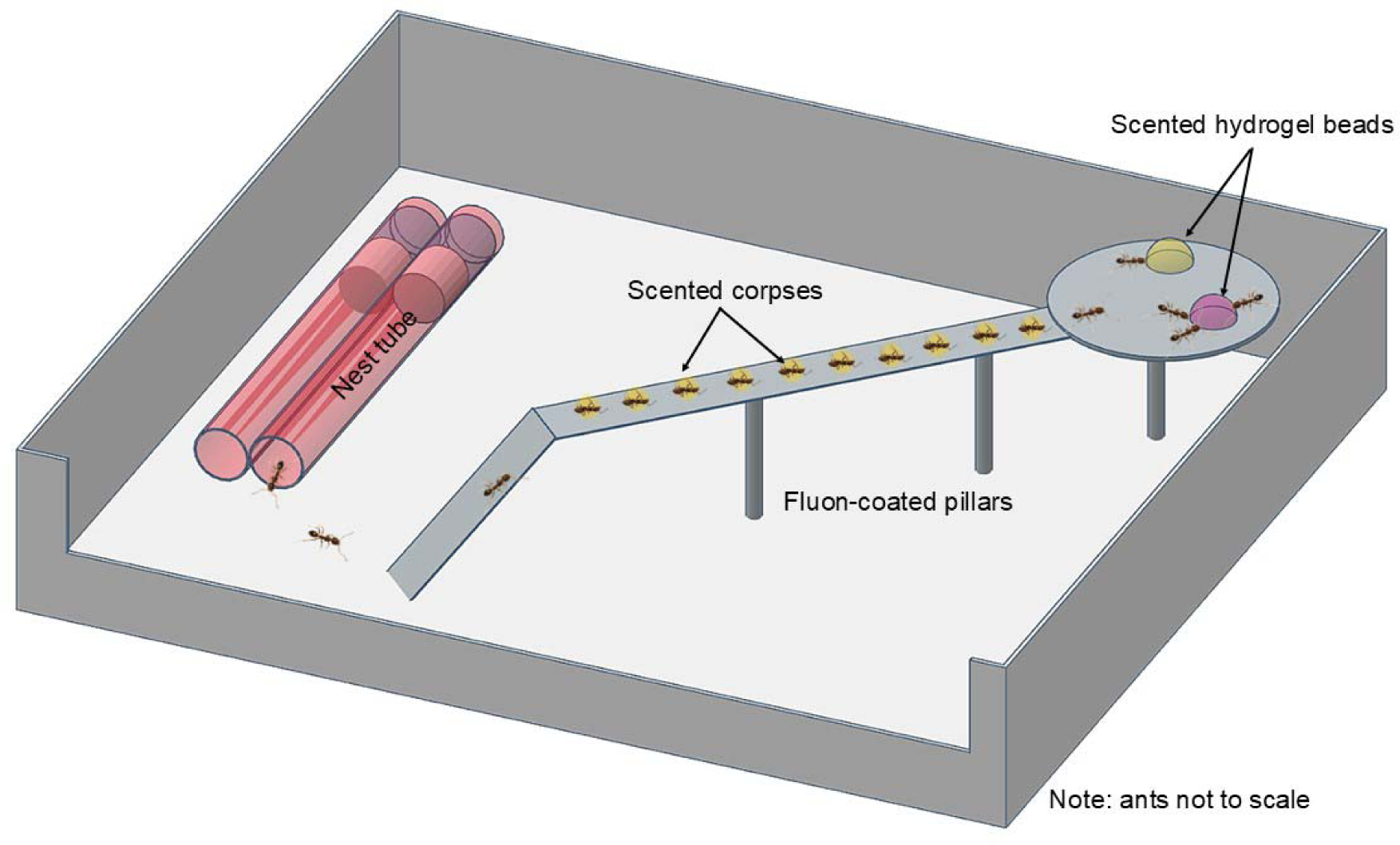
Collective decision-making experiment. A round plastic platform, supported by fluon-coated pillars and reachable by a bridge, was placed in the colony box. Hydrogel beads soaked in flavoured sucrose solution (apple and strawberry) were offered. 12 scented nestmate corpses were placed on the bridge. Foraging ants had to pass the scented corpses to reach the feeding platform. The number of ants within 0.5 cm of each food type were counted.

### Statistical analysis

Data were analysed using generalized linear mixed-effect models (GLMM) (34) in R version 4.1.0 (R Core Team 2021). GLMMs were fitted using the lme4 package (35). Because the data were binomial in experiments 1 & 2 (novel/corpse odour choice) and in experiment 3 (proportion drinking time on feeder A vs feeder B), a binomial error distribution was used. For experiment 4, a Zero-Inflated Beta Model (for proportion data) and a glmm with a poisson error distribution (for ant counts on different feeders over time) was used. Because multiple ants were tested per colony, we included colony as random factor. The R package emmeans was used for pairwise contrasts. It allows for the estimation and comparison of means across different levels of categorical predictors in a model. Model fit was checked using the DHARMa package (36). Results were plotted using the gglot2 package (37). The complete code and analysis output is provided in supplements 1.

Note that in our analysis, we excluded four very small nests from the models (40 ants or less foraging over 24 hours). For statistical information on all colonies, see supplement 2. Therein we also provide a complete analysis of the dataset including the four excluded colonies. The results of that analysis show the same patterns as those presented here. We also analyse the data in terms of proportion of ants on each food type, rather than raw count. Again, the results are similar, and are presented in supplement 2.

## Results

### Experiment 1 – Testing the effect of corpse-scent association on odour preference using scented corpses

Ants that had encountered scented corpses chose the Y-maze odour arm associated with the corpses significantly less often than the opposite scented arm (69% chose the novel odour arm, GLMM, z-ratio = 2.92, p = 0.003, see fig. 3A). By contrast, ants in the control group that experienced the odour associated with scented dummies showed no significant preference between the two scented arms (47% chose the novel-odour arm, GLMM, z-ratio = -0.5, p = 0.62, see fig.3A). The treatment groups differed significantly from each other (emmeans, z-ratio = 2.48, p = 0.013, see fig. 3A). The specific corpse/dummy odour (blackcurrent or orange) did not significantly affect choice (GLMM: z-ratio = 0.37, p = 0.71).

**Figure 3:**
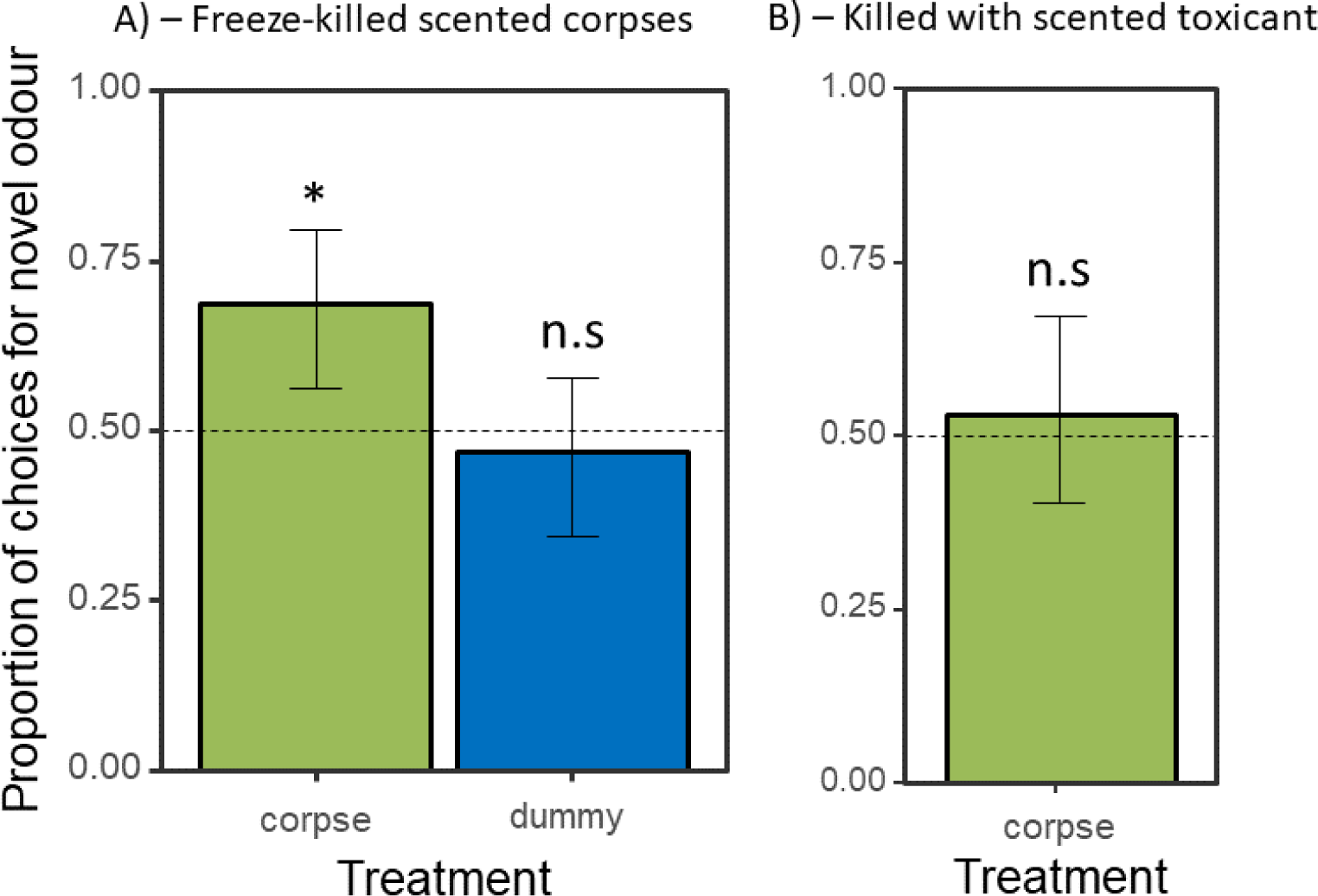
The response of ants to odours associated with nestmate corpses (experiments 1 & 2): A) Experiment 1: Proportion of choices for novel odour, per treatment group, corpses or dummies (control). B) Proportion of ants which encountered nestmate corpses killed by feeding on scented food choosing the novel scented Y-maze arm. Note that since experiment 2 relied on ants consuming the scented, toxic bait, no control was possible. Bars depict means, whiskers 95% confidence intervals derived from the fitted GLMM. The dotted horizontal line displays chance level of 50%. *_p_ < 0.05; **_p_ < 0.01; ***p < 0.001.

### Experiment 2 – Testing the effect of corpse-scent association on odour preference using ants killed with scented toxicant

The ants showed no difference in their choice of odour arm (GLMM, z-ratio = 0.58, p = 0.56, fig.3B): 53% of the ants that were exposed to corpses fed with a flavoured toxicant chose the arm scented with the other odour. The specific odour associated with the corpses did not significantly affect arm choice (GLMM: z-ratio = -0.51, p = 0.61). Unexpectedly, focal ants were commonly observed to lick the mouths of the corpses.

### Experiment 3 – Testing the effect of corpse-associated odour on feeding preference

Ants starved for two days did not feed significantly longer on either of the food sources, irrespective of their treatment (GLMM, dummy: z-ratio = 1.74, p = 0.08, 60% fed longer from the novel odour. Corpse: z-ratio = 0.51, p = 0.61, 53% fed for longer from the novel feeder, see fig. 4A). Odour did not significantly affect these measures (GLMM, feeding time, z-ratio = 0.55, p = 0.58). Similar outcomes were found for ants starved for 4 days feed (GLMM, dummy, z-ratio = -0.56, p = 0.58, 42% fed longer from the novel feeder, corpse, z-ratio = 0.87, p = 0.39, 55% fed longer from the novel feeder, see fig. 4B). Odour type did not significantly affect choice (GLMM, feeding time, z-ratio = -0.25, p = 0.80).

**Figure 4:**
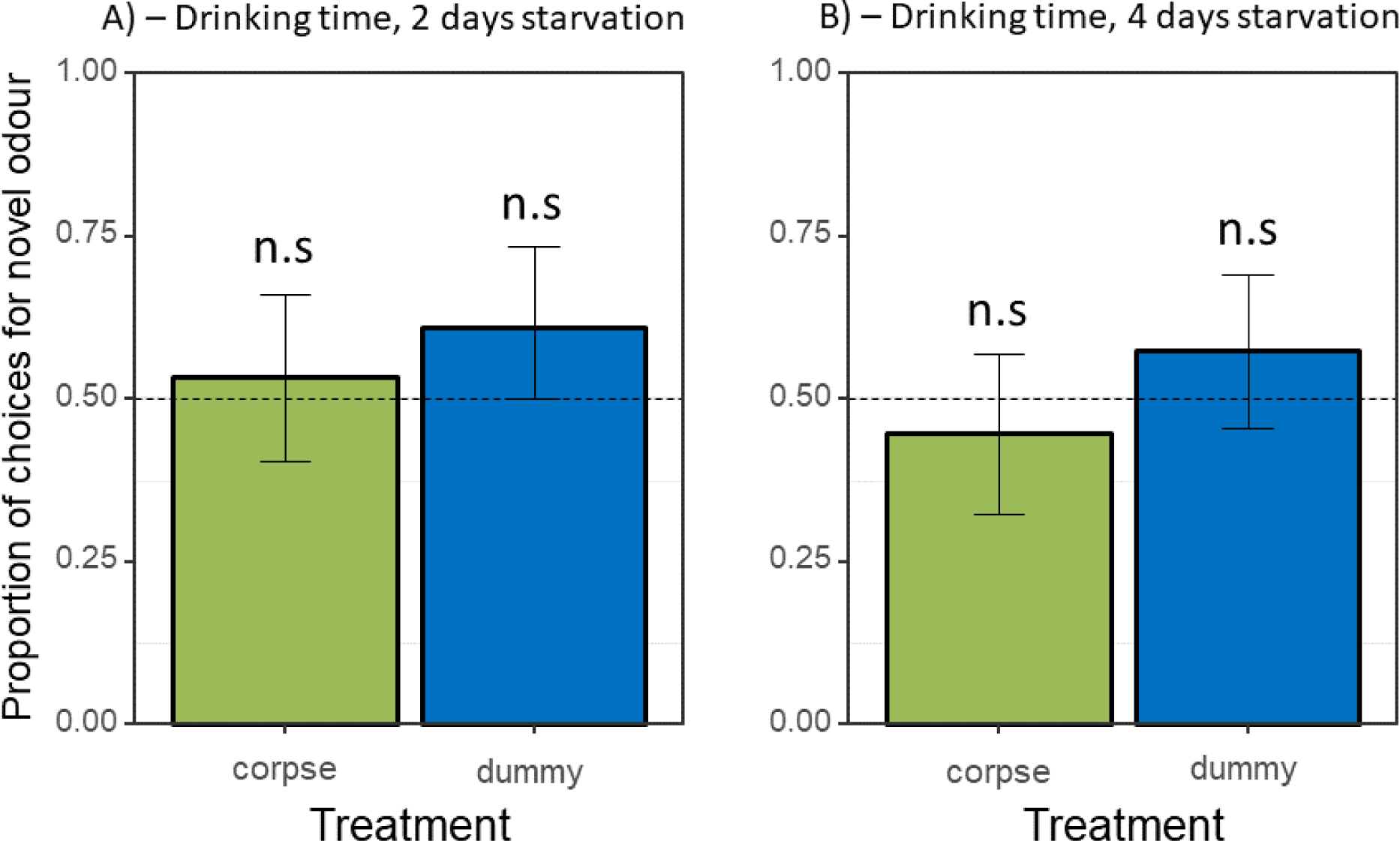
Effect of corpse-associated smell on feeding preference (exp 3): A) and B) show the mean (±95% confidence intervals) proportion of time spent feeding at the feeder not scented to match the corpses or dummies, by ants starved for 2 and 4 days respectively.

### Experiment 4 – Can the presence of scented corpses drive collective food choice?

In the first 60 minutes significantly fewer ants (705/1769) fed on the hydrogel beads scented to match the corpses (emmeans: t-ratio = -7.751 p < 0.0001, fig. 5A/B). Over the following 23h, foraging levels were extremely low (mean 0.324 ants on each feeder at each timepoint per colony), with no clear preference for the novel feeder (67/111 ants at the novel feeder, emmeans: t-ratio = -1.163, p = 0.2456, fig. 5C/D. The specific corpse odour did not significantly affect their choice within 60 minute (GLMM: z-ratio = -5.407, p = 0.8669) nor the following 23 hours (GLMM: z-ratio = 1.203, p = 0.2291).

**Figure 5:**
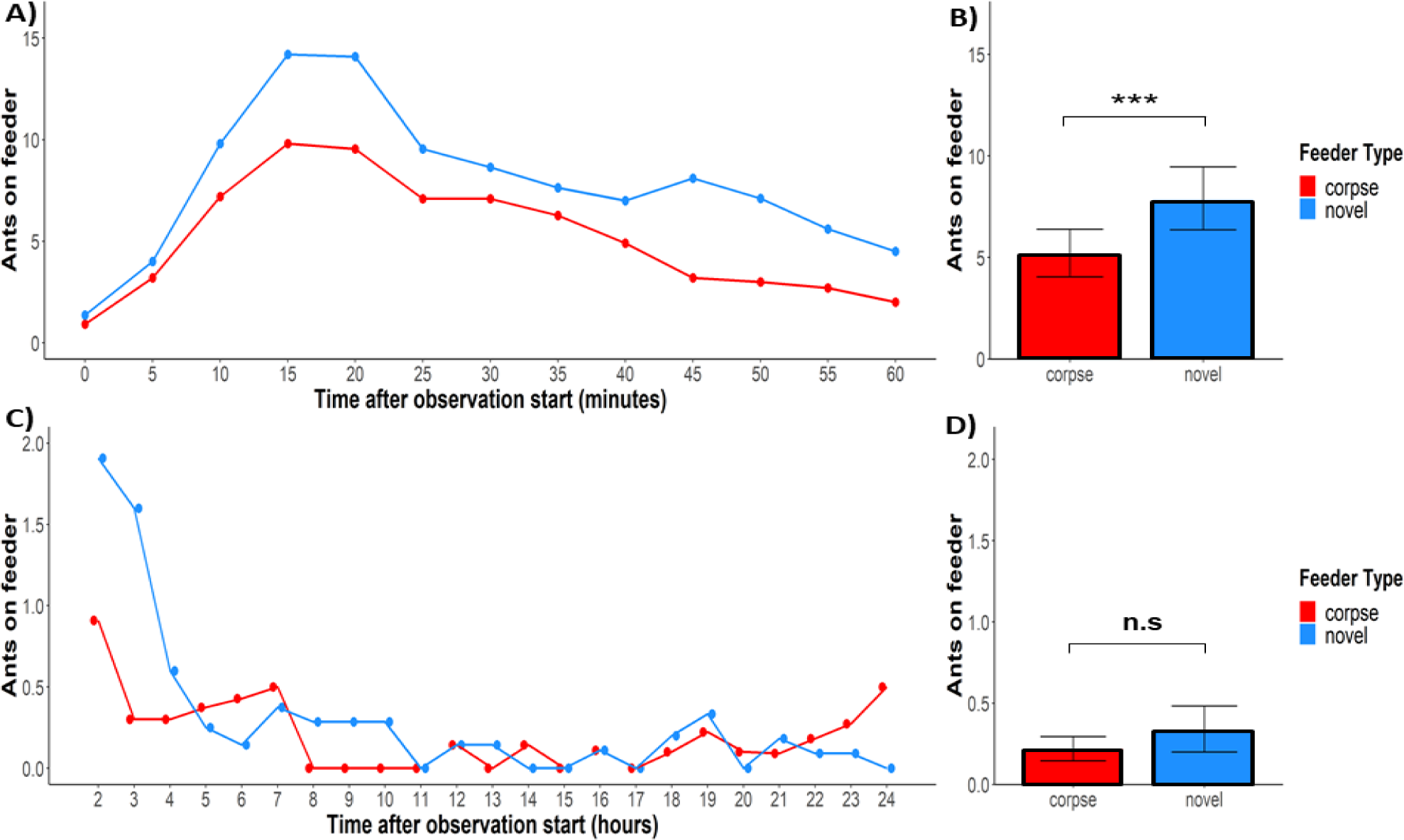
Collective foraging effort on feeders scented to match either nestmate-corpses encountered en-route to the feeder, or scented with a novel odour (exp 4): A) Dual-curve plot, showing average number of ants within 1 cm of feeder, at 5 minute intervals for 60 minutes post food presentation. B) Bar plot, Mean (±SE) number of ants sticking 1 cm or closer to a feeder over the first 60 minutes. C/D) corresponding to A/B but 23 hours post food presentation in 1 hour intervals. Bars depict means, whiskers 95% confidence intervals derived from the fitted GLMM or Zero-Inflated Beta Model. N counted ants (60min) = 1769, N counted ants (23hours) =111. A total of 12 trials were conducted.

## Discussion

Our study strongly suggests that nestmate corpses are a negative stimulus for the ant *Linepithema humile*. The implication is that corpses may drive toxic bait avoidance at an individual and collective level, by causing the ants to avoid odours associated with them (experiments 1 and 4). However, under conditions more relevant to ant control efforts – the consumption of scented toxicant-laced food (experiment 2) no avoidance was found. Similarly, in the dual-feeder experiment (experiment 3), ants did not feed less from the solution scented to match the odour of scented corpses.

Avoidance of the scented Y-maze arms associated with nestmate corpses (experiment 1) demonstrates that corpses can act as a negative stimulus for *L. humile*. Avoidance of cues associated with negative consequences has been demonstrated in ants previously, with ants avoiding Y-maze or T-maze arms scented with a bitter quinine solution (29,38). Nestmate corpses have also been shown to be avoided by ants, even to the point that potential nest sites containing foreign ant corpses were avoided by Temnothorax albipennis (21). However, we note that ants can discriminate between foreign or nestmate corpses (22). It is thus unclear if the reason for avoidance in the nest site avoidance study was to avoid a confrontation with a foreign ant colony, or for hygienic reasons, or both. However, given that all the ants in our study stemmed from the same supercolony, and showed no avoidance or aggression between colony fragments, it is unlikely that non-nestmate avoidance plays a large role in the results. Although our study showed a relatively high avoidance rate (69%), it is not clear why avoidance was not higher, such as for avoidance of nests containing corpses (21). We note that a certain willingness to approach nestmate corpses is required, in order for the corpses to be cleared away, and indeed observed several ants handling, moving, and even beginning to dismember corpses. Corpse clearance has been shown to be an important hygienic response, preventing disease spread (18). It was even shown in Argentine ants that some workers have to apply regular pygidial gland secretions on corpses to prevent growth of pathogenic fungi (39), which requires the corpses to be approached.

Surprisingly, our second experiment, which had the same setup as the first experiment, found no avoidance of the corpse-associated odour. The key difference between these experiments was how the ants that were used as corpse stimuli, died and were stored. In the first experiment the ants were freeze-killed then stored for 24h in a Petri dish containing food flavouring, whereas the ants in the second experiment were allowed to feed on toxic, flavoured hydrogel beads, subsequently died within 8h, and were then stored for a further 16h in a Petri dish without any food flavour. It is possible that in this second method the test ants might have not perceived the flavour of the poisoned food on the corpses, because it was too subtle or disappeared after 24 hours. Ants can identify nestmates as corpses within 1 hour of death (20), so toxicants with immediate lethal effects may be more detectible, because bait odours may linger long enough on the corpses, whereas toxicants at levels which cause delayed action may be less detectible. We also note that many commercial baits are composed of a matrix with a potentially longer-lasting odour profile e.g. fat or protein-rich pastes (10,40–42). It is possible that such baits imbue ants which fed on them with a long-lasting odour that could drive future rejection when these ants finally die. The water-soluble odour we used in the current study may not have lasted long enough to be detectable by other ants 16-24 hours after feeding.

Interestingly, we observed that most test ants licked the mouths of the dead ants, possibly tasting the food consumed by the poisoned nestmates. They thus might have formed a direct positive association between the odour of the food and it’s sweetness, which may have interfered with, or counterbalanced, any negative stimulus created by the corpses. Ants have been demonstrated to learn food-related odour cues from returning foragers, even without active food sharing (14–16,43). The results of this experiment thus do not rule out that corpses act as a negative stimulus in these ants, but imply that perhaps this effect will not strongly drive behaviour in a real-world ant control situation using sucrose-based baits. However, other commonly-used bait types based on fat or protein pastes may have elicited an effect (see below).

The third experiment showed that ants did not feed less from a food source flavoured to match the scented corpses than from a neighbouring food source offering a novel flavour, irrespective of whether the ants were starved for 2 or 4 days. It is unlikely that the ants were unable to associate the odour impregnated onto the corpses with the taste of the flavoured sucrose feeder because we have previously (26) demonstrated that Argentine ants can learn to associate a flavoured food reward with the corresponding environmental odour after only one exposure. Further experiments with this assay offering 1M vs 0.5M sucrose, or 1M vs 1M sucrose + a low level of bitter quinine, have shown that this method is effective at uncovering food preference (unpublished data). A more likely reason for this unexpected result was the overwhelming drive to feed under even moderate starvation conditions: The ants simply continued feeding at the first food source they encountered, and any potential negative stimulus associated with the food flavour was not sufficiently strong to override ongoing feeding. Individual *L. humile* workers are unusually motivated feeders (44). Alternatively, ants may only show avoidance in terms of orientation, not food rejection once the food is encountered. In other words, the ants might only avoid going towards areas scented to match the scent of corpses, but do not avoid feeding at similarly-flavoured food sources.

The fourth experiment tested colony-level preference, by examining collective foraging choices. Collectively, over the first hour of foraging where the vast majority of foraging occurred, the ants avoided the corpse-associated hydrogel beads and preferred to feed on the alternative beads. The proximate mechanism driving this pattern is unclear: ants may have avoided approaching the corpse-associated bead, may have left the bead before feeding fully, may have down-modulated their recruitment to the bead, or any combination of these options. Unfortunately, we lack detailed information on the behaviour of the individual ants in the collective foraging assay, especially the first few foragers, which would be needed to distinguish these mechanisms. Observations over the first 10 minutes suggested that ants accepted the first bead they contacted, regardless of whether it was the corpse- or control bead. This fits well with the results from experiment 3, and makes at least the taste-and-reject hypothesis unlikely. Due to the positive feedback inherent in ant mass-recruitment systems, the behaviour of the first ants in such experiments has a decisive influence on the final collective outcome (45). Thus, if the first few ants tended to preferentially approach the novel bead, and laid the first pheromone trail in that direction, this would bias all future choices towards that bead. This effect would compound the tendency of the ants to avoid approaching the corpse-associated odour.

Finally, a second positive-feedback process may well be in place: The first ants to feed may have encountered naïve outgoing ants on their return journey, and certainly shared food via trophallaxis with other ants in the nest. Such on-trail and within-nest encounters could have created a positive association to the smell of the bead area, causing these newly recruited ants to preferentially choose that area – as has already been shown in this and other species (14,15,26). Note that in systems characterised by strong positive feedback, such as this one, stochasticity can play a very large role. It is thus not surprising that in a subset of experiments the corpse-associated bead was chosen, as even weaker options, if discovered first, can rapidly outcompete stronger options discovered later (33,46). Our study demonstrates that *Linepithema humile* can form negative associations with a scent associated with nestmate corpses, which leads them to avoid areas with the corresponding odour. However, this aversion may not be strong enough to override positive stimuli, such as the presence of a desired food. These findings have important implications for ant control strategies, as they suggest that the presence of nestmate corpses may reduce bait efficacy. The formation of a negative association between a bait and affected nestmates may negatively impact the efficacy of control efforts. However, given the lack of effect found in experiment 2, which used more field-realistic methods, it is unclear whether this effect plays a major role in real-world control efforts. Nevertheless, if this association-driven avoidance of baits is indeed impacting control efforts, a straightforward intervention suggests itself: baits can be scented with distinct odours, and odours can be either cycled between treatments, or multiple odours can be deployed simultaneously. Both interventions should reduce the ability of ants to form firm associations between the bait odour and the odour of killed ants. This approach may well have other benefits as well, such as making the baits easier for ants to find. The commercial availability of cheap and environmentally benign food flavourings makes such an intervention easily and broadly deployable.

## Supporting information

Supplement 2 Experiment 4 All Colonies and Proportion Statistics

Supplement 1 Complete R Studio Markdown

## Declaration of Interest

None.

## Author contribution

T. Wagner and T. J. Czaczkes designed the study. T. Wagner collected the data. T. Wagner analysed the data and wrote the manuscript. T. J. Czaczkes edited the manuscript. All authors read and approved the manuscript.

## Acknowledgements

We would like to thank Silvia Abril and Eduardo Sequeira for providing ant colonies, and Roxana Josens and Ben Hoffmann for advice on the manuscript. Thomas Wagner was supported by an ERC starter grant (Cognitive control: 948181) to Tomer J. Czaczkes. Tomer J. Czaczkes was supported by a Heisenberg fellowship from the Deutsche Forschungsgemeinschaft (CZ 237 / 4-1).

## Notes

### Competing Interest Statement

The authors have declared no competing interest.

### Summary of Updates

Minor text improvements and figure 4 x-axis correction.

