## Supplement 2 Experiment 4 All Colonies and Proportion Statistics for "Food odours associated with conspecific corpses cause food avoidance in an invasive ant"

This file provides the statistical analysis summary of experiment 4 including the two very small colonies. For the complete statistical analysis output, see supplement 1.

### Experiment 4 – Can the presence of scented corpses drive collective food choice (all colonies)?

In the first 60 minutes significantly fewer ants (784/1881) fed on the hydrogel beads scented to match the corpses (emmeans: t-ratio = -7.440, p < 0.0001, fig. S1A/B). Over the following 23 hours, ants did not show a trend towards avoiding the feeder (78/127) associated with the corpses (emmeans: t-ratio = -1.553, p = 0.1210, fig. S1C/D). The specific corpse odour did not significantly affect their choice within 60 minutes (GLMM: z-ratio = 0.168, p = 0.8669) nor the following 23 hours (GLMM: z-ratio = 1,237, p = 0.216). The proportion of ants feeding on hydrogel beads scented to match the corpses was significantly lower after 60 minutes than on the novel feeder (emmeans: t-ratio = -2.777, p < 0.0055, fig. S1C). Over the following 23 hours, the proportion of ants feeding on the novel feeder, was not significantly higher than on the corpse associated feeder (emmeans: t-ratio = -1.508, p = 0.1317, fig. S1F).

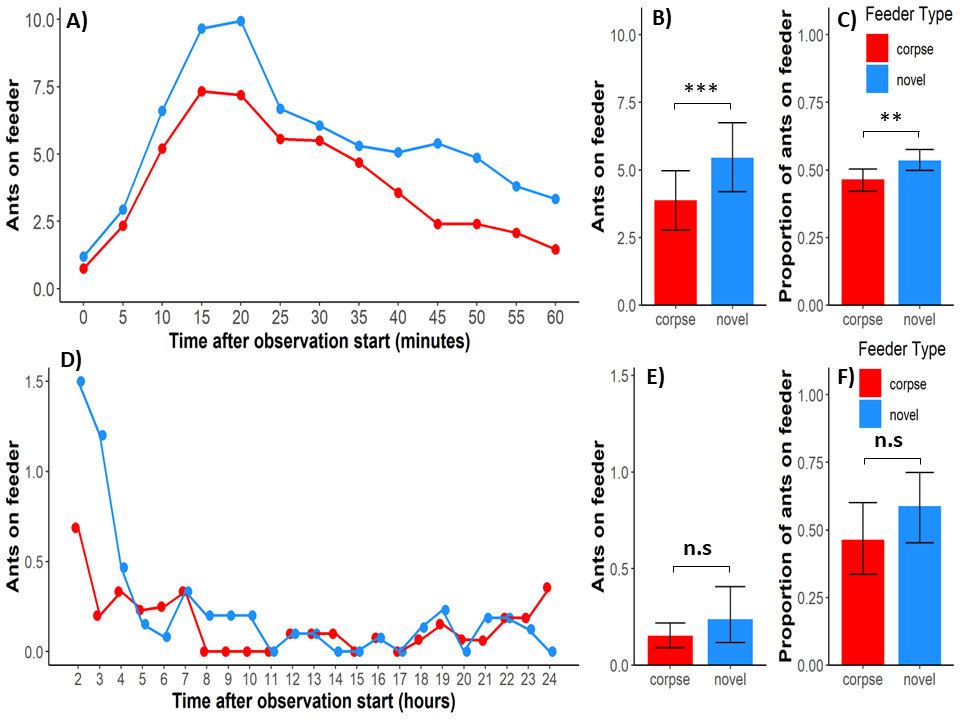

**Figure S1:** Collective foraging effort on feeders scented to match either nestmate-corpses encountered en-route to the feeder, or scented with a novel odour (exp 4): A) Dual-curve plot, average number of ants sticking 1 cm or closer to a feeder, 60 minutes post food presentation per 5 minutes intervals. B) Bar plot, average number of ants sticking 1 cm or closer to a feeder over the first 60 minutes. C) Bar plot, proportion of ants sticking 1cm or closer to a feeder over the first 60 minutes. D/E/F corresponding to A/B/C but 23 hours post food presentation in 1 hour intervals. Bars depict means, whiskers 95% confidence intervals derived from the fitted GLMM or Zero-Inflated Beta Model. Note the different scales of figures A/B/C and D/E/F. N counted ants (60min) = 1881, N counted ants (23hours) = 127. All in all 12 trials were conducted.

### Experiment 4 – Can the presence of scented corpses drive collective food choice (without small colonies, only proportion)?

The proportion of ants feeding on hydrogel beads scented to match the corpses was significantly lower after 60 minutes than on the novel feeder (emmeans: t-ratio = -2.907, p < 0.0036, fig. S2A). Over the following 23 hours, the proportion of ants feeding on the novel feeder, was not significantly higher than on the corpse associated feeder (emmeans: t-ratio = -1.519, p = 0.1288, fig. S2B).

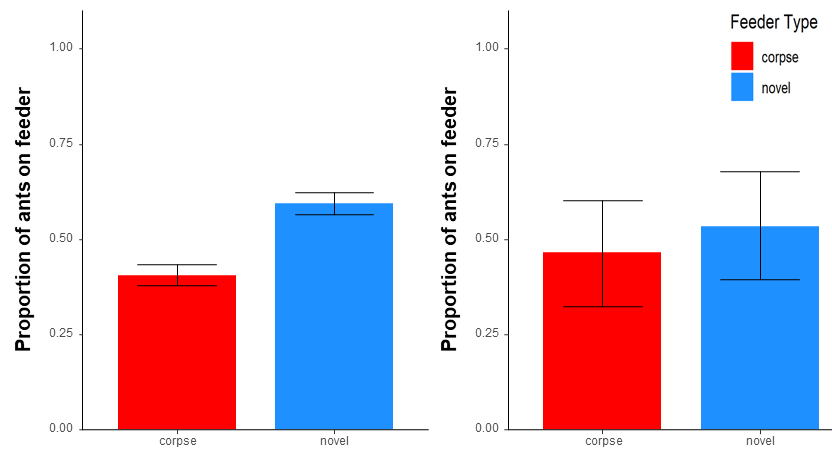

**Figure S2:** Collective foraging effort on feeders scented to match either nestmate-corpses encountered en-route to the feeder, or scented with a novel odour (exp 4): A) Bar plot, proportion of ants sticking 1cm or closer to a feeder over the first 60 minutes. B) corresponding to A) but 23 hours post food presentation in 1 hour intervals. Bars depict means, whiskers 95% confidence intervals derived from the fitted GLMM or Zero-Inflated Beta Model. Note the different scales of figures A and B. N counted ants (60min) = 1769, N counted ants (23hours) = 111. All in all 12 trials were conducted.
