## Supplement 1 Complete R Studio Markdown for "Food odours associated with conspecific corpses cause food avoidance in an invasive ant"

2023-07-20

### Data import, libraries, housekeeping

loading libraries

library (ggplot2)
library (gplots)
library (ggpubr)
library (lme4)
library (nlme)
library (tidyverse)
library (emmeans)
library (multcomp)
library (DHARMa)
library (readxl)
library (wesanderson)
library (grid)
library (gridExtra)
library (dplyr)
library (scales)
library (lattice)
library (MASS)
library(EnvStats)
library(officer)
library(rvg)
library(Hmisc)
library(eoffice)
library(cowplot)
library (glmmTMB)
require(pscl) # alternatively can use package ZIM for zero-inflated models
library(lmtest)
library(WebPower)
library(ggthemes)
library(forcats)
library(stringr)
library(simr)
library(betareg)
library(pscl)
library(gamlss)
library(extrafont)

setting directory

import all data files for later analysis.

#### 1. Analysis, Experiment 1 – Scented corpses, runway odour on a Y-maze

##### Glmm modeling, pairwise, Experiment 1 – Testing the effect of corpse-scent association on odour preference using scented corpses

```
#Glmm#

Exp1negative$treatment<-as.factor(Exp1negative$treatment)

Exp1negative_m <-glmer(correct_final ~ treatment + dead_odour + (1|colony),
                                family="binomial",
                                data= Exp1negative)
## boundary (singular) fit: see help('isSingular')
summary(Exp1negative_m)
## Generalized linear mixed model fit by maximum likelihood (Laplace
##   Approximation) [glmerMod]
##  Family: binomial  ( logit )
## Formula: correct_final ~ treatment + dead_odour + (1 | colony)
##    Data: Exp1negative
##
##      AIC      BIC   logLik deviance df.resid
##    175.8    187.2    -83.9    167.8      124
##
## Scaled residuals:
##     Min      1Q  Median      3Q     Max
## -1.5347 -0.9715  0.6516  0.6970  1.1012
##
## Random effects:
##  Groups Name        Variance Std.Dev.
##  colony (Intercept) 0        0
## Number of obs: 128, groups:  colony, 4
##
## Fixed effects:
##                  Estimate Std. Error z value Pr(>|z|)
## (Intercept)        0.7219     0.3238   2.230   0.0258 *
## treatmentdummy    -0.9146     0.3683  -2.483   0.0130 *
## dead_odourOrange   0.1348     0.3673   0.367   0.7136
## ---
## Signif. codes:  0 '***' 0.001 '**' 0.01 '*' 0.05 '.' 0.1 ' ' 1
##
## Correlation of Fixed Effects:
##             (Intr) trtmnt
## tretmntdmmy -0.602
## dead_drOrng -0.553 -0.015
## optimizer (Nelder_Mead) convergence code: 0 (OK)
## boundary (singular) fit: see help('isSingular')
#Pairwise#

Pairwise1 = emmeans(Exp1negative_m, spec= "treatment")

Pairwise1=contrast(Pairwise1, method = "pairwise")
summary(Pairwise1)
##  contrast       estimate    SE  df z.ratio p.value
##  corpse - dummy    0.915 0.368 Inf   2.483  0.0130
##
## Results are averaged over the levels of: dead_odour
## Results are given on the log odds ratio (not the response) scale.
#Separate treatment#

Exp1negative_dummy <- subset (Exp1negative, treatment=="dummy")
Exp1negative_corpse <- subset (Exp1negative, treatment=="corpse")


Exp1negative_dummy_m <-glmer(correct_final ~ 1 + (1|colony),
                                             family="binomial",
                                             data= Exp1negative_dummy )
## boundary (singular) fit: see help('isSingular')
summary(Exp1negative_dummy_m)
## Generalized linear mixed model fit by maximum likelihood (Laplace
##   Approximation) [glmerMod]
##  Family: binomial  ( logit )
## Formula: correct_final ~ 1 + (1 | colony)
##    Data: Exp1negative_dummy
##
##      AIC      BIC   logLik deviance df.resid
##     92.5     96.8    -44.2     88.5       62
##
## Scaled residuals:
##     Min      1Q  Median      3Q     Max
## -0.9393 -0.9393 -0.9393  1.0646  1.0646
##
## Random effects:
##  Groups Name        Variance Std.Dev.
##  colony (Intercept) 0        0
## Number of obs: 64, groups:  colony, 4
##
## Fixed effects:
##             Estimate Std. Error z value Pr(>|z|)
## (Intercept)  -0.1252     0.2505    -0.5    0.617
## optimizer (Nelder_Mead) convergence code: 0 (OK)
## boundary (singular) fit: see help('isSingular')
Exp1negative_corpse_m <-glmer(correct_final ~ 1 + (1|colony),
                                             family="binomial",
                                             data= Exp1negative_corpse )
## boundary (singular) fit: see help('isSingular')
summary(Exp1negative_corpse_m)
## Generalized linear mixed model fit by maximum likelihood (Laplace
##   Approximation) [glmerMod]
##  Family: binomial  ( logit )
## Formula: correct_final ~ 1 + (1 | colony)
##    Data: Exp1negative_corpse
##
##      AIC      BIC   logLik deviance df.resid
##     83.5     87.8    -39.7     79.5       62
##
## Scaled residuals:
##     Min      1Q  Median      3Q     Max
## -1.4832 -1.4832  0.6742  0.6742  0.6742
##
## Random effects:
##  Groups Name        Variance Std.Dev.
##  colony (Intercept) 0        0
## Number of obs: 64, groups:  colony, 4
##
## Fixed effects:
##             Estimate Std. Error z value Pr(>|z|)
## (Intercept)   0.7885     0.2697   2.924  0.00346 **
## ---
## Signif. codes:  0 '***' 0.001 '**' 0.01 '*' 0.05 '.' 0.1 ' ' 1
## optimizer (Nelder_Mead) convergence code: 0 (OK)
## boundary (singular) fit: see help('isSingular')
```

- Ants experiencing corpses were significantly different in their final choice than dummies (pairwise)
- While ants experiencing dummies were not choosing a side/odour significantly more often, while the ants experiencing corpses chose significantly more often the side/odour contrary to the corpses odour.

Does this main model work?

dharmit1 <-simulateResiduals(Exp1negative_m)
plot(dharmit1)

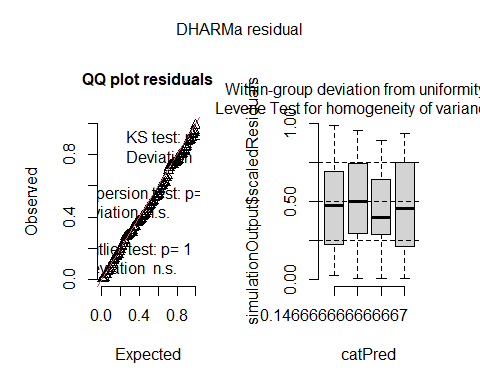
 - Yes it works.

##### Figures

```
fig1N <- ggplot (Exp1negative, aes(x = treatment, y = correct_final))+
  scale_y_continuous(expand = c(0, 0)) + # forces X axis to 0, but in this case is overriden by ribbon
  geom_point( alpha = 0) +
  stat_summary(fun.y = "mean", geom = "bar", fill="dodgerblue4") +
  ylab("Proportion of choices for non-corpse odour") +
  xlab("Treatment") + stat_summary(fun.data = "mean_cl_boot", geom="errorbar", width = 0.2) +
  theme_bw(20)+
  geom_abline(intercept = 0.5, slope = 0, color = "black", linetype = 2) +
  coord_cartesian( ylim = c(0, 1.0)) # zooms in on the top half
## Warning: `fun.y` is deprecated. Use `fun` instead.
print(fig1N)
```

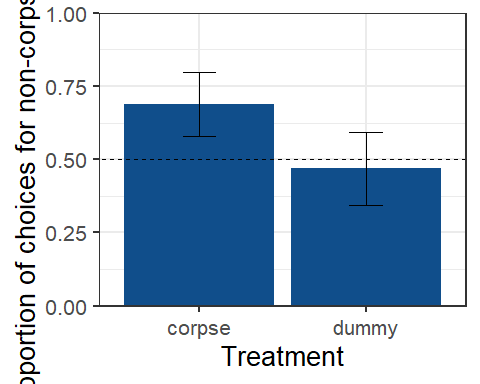

#### 2. Analysis, Experiment 2 – Scented corpses, runway odour on a Y-maze

##### Experiment 2 – Testing the effect of corpse-scent association on odour preference using corpses killed with scented poison

```
#Glmm#

Exp2negative$dead_flavour<-as.factor(Exp2negative$dead_flavour)

Exp2negative_m <-glmer(dead_final_avoidance ~ dead_flavour + dead_side + (1|colony),
                                family="binomial",
                                data= Exp2negative)
## boundary (singular) fit: see help('isSingular')
summary(Exp2negative_m)
## Generalized linear mixed model fit by maximum likelihood (Laplace
##   Approximation) [glmerMod]
##  Family: binomial  ( logit )
## Formula: dead_final_avoidance ~ dead_flavour + dead_side + (1 | colony)
##    Data: Exp2negative
##
##      AIC      BIC   logLik deviance df.resid
##     95.2    103.8    -43.6     87.2       60
##
## Scaled residuals:
##     Min      1Q  Median      3Q     Max
## -1.2896 -1.0010  0.7754  0.8810  1.1350
##
## Random effects:
##  Groups Name        Variance Std.Dev.
##  colony (Intercept) 0        0
## Number of obs: 64, groups:  colony, 4
##
## Fixed effects:
##                    Estimate Std. Error z value Pr(>|z|)
## (Intercept)          0.2533     0.4371   0.580    0.562
## dead_flavourOrange   0.2553     0.5060   0.505    0.614
## dead_sideR          -0.5067     0.5061  -1.001    0.317
##
## Correlation of Fixed Effects:
##             (Intr) dd_flO
## dd_flvrOrng -0.565
## dead_sideR  -0.579 -0.016
## optimizer (Nelder_Mead) convergence code: 0 (OK)
## boundary (singular) fit: see help('isSingular')
#Pairwise#
Pairwise2 = emmeans(Exp2negative_m, spec= "dead_flavour")

Pairwise2=contrast(Pairwise2, method = "pairwise")
summary(Pairwise2)
##  contrast        estimate    SE  df z.ratio p.value
##  Cassis - Orange   -0.255 0.506 Inf  -0.505  0.6138
##
## Results are averaged over the levels of: dead_side
## Results are given on the log odds ratio (not the response) scale.
#Separate flavour#

Exp2negative_orange <- subset (Exp2negative, dead_flavour=="Orange")
Exp2negative_cassis <- subset (Exp2negative, dead_flavour=="Cassis")


Exp2negative_orange_m <-glmer(dead_final_avoidance ~ 1 + (1|colony),
                                             family="binomial",
                                             data= Exp2negative_orange )
## boundary (singular) fit: see help('isSingular')
summary(Exp2negative_orange_m)
## Generalized linear mixed model fit by maximum likelihood (Laplace
##   Approximation) [glmerMod]
##  Family: binomial  ( logit )
## Formula: dead_final_avoidance ~ 1 + (1 | colony)
##    Data: Exp2negative_orange
##
##      AIC      BIC   logLik deviance df.resid
##     47.9     50.8    -21.9     43.9       30
##
## Scaled residuals:
##     Min      1Q  Median      3Q     Max
## -1.1339 -1.1339  0.8819  0.8819  0.8819
##
## Random effects:
##  Groups Name        Variance Std.Dev.
##  colony (Intercept) 0        0
## Number of obs: 32, groups:  colony, 2
##
## Fixed effects:
##             Estimate Std. Error z value Pr(>|z|)
## (Intercept)   0.2513     0.3563   0.705    0.481
## optimizer (Nelder_Mead) convergence code: 0 (OK)
## boundary (singular) fit: see help('isSingular')
Exp2negative_cassis_m <-glmer(dead_final_avoidance ~ 1 + (1|colony),
                                             family="binomial",
                                             data= Exp2negative_cassis )
## boundary (singular) fit: see help('isSingular')
summary(Exp2negative_cassis_m)
## Generalized linear mixed model fit by maximum likelihood (Laplace
##   Approximation) [glmerMod]
##  Family: binomial  ( logit )
## Formula: dead_final_avoidance ~ 1 + (1 | colony)
##    Data: Exp2negative_cassis
##
##      AIC      BIC   logLik deviance df.resid
##     48.4     51.3    -22.2     44.4       30
##
## Scaled residuals:
##    Min     1Q Median     3Q    Max
##     -1     -1      0      1      1
##
## Random effects:
##  Groups Name        Variance Std.Dev.
##  colony (Intercept) 0        0
## Number of obs: 32, groups:  colony, 2
##
## Fixed effects:
##               Estimate Std. Error z value Pr(>|z|)
## (Intercept) -3.053e-16  3.536e-01       0        1
## optimizer (Nelder_Mead) convergence code: 0 (OK)
## boundary (singular) fit: see help('isSingular')
```

- Ants did not show any significantly aversion for the odour associated to the corpse odour smell.

Does this model work?

dharmit2 <-simulateResiduals(Exp2negative_m)
plot(dharmit2)

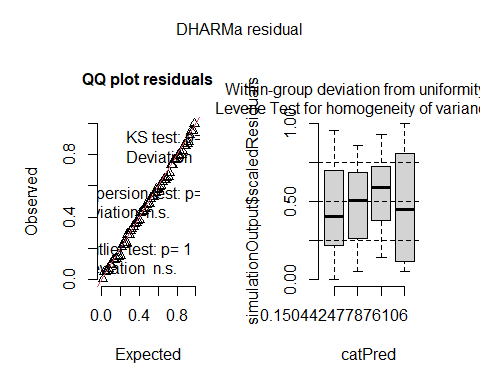
 - Yes it works.

##### Figures Note figures were later merged and partly edited in powerpoint

```
fig2N <- ggplot (Exp2Merged, aes(x = Treatment, y = dead_final_avoidance ))+
  scale_y_continuous(expand = c(0, 0)) + # forces X axis to 0, but in this case is overriden by ribbon
  geom_point( alpha = 0) +
  stat_summary(fun.y = "mean", geom = "bar", fill="dodgerblue4") +
  ylab("Proportion of choices for non-corpse odour") +
  xlab("Treatment") + stat_summary(fun.data = "mean_cl_boot", geom="errorbar", width = 0.2) +
  theme_bw(18) +
  geom_abline(intercept = 0.5, slope = 0, color = "black", linetype = 2) +
  coord_cartesian( ylim = c(0, 1.0)) # zooms in on the top half
## Warning: `fun.y` is deprecated. Use `fun` instead.
print(fig2N)
```

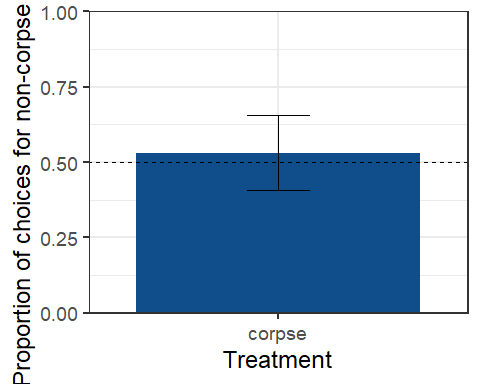

#### 3. Analysis, Experiment 3 – Scented corpses, food flavour on a two-choice delta feeder, 2 days and 4 days starvation

##### Glmm modeling, pairwise, Experiment 3 – Testing the effect of corpse-associated odour on feeding preference

```
# 2 days starvation


# GLMM drinking_time

Exp3bnegative_m <-glmmTMB(Drinking_aversive ~ corpse_dummy + dead_odour + (1|colony),
                              family="binomial",
                              data= Exp3negative)
summary(Exp3bnegative_m)
##  Family: binomial  ( logit )
## Formula:          Drinking_aversive ~ corpse_dummy + dead_odour + (1 | colony)
## Data: Exp3negative
##
##      AIC      BIC   logLik deviance df.resid
##    179.0    190.4    -85.5    171.0      122
##
## Random effects:
##
## Conditional model:
##  Groups Name        Variance  Std.Dev.
##  colony (Intercept) 7.532e-10 2.744e-05
## Number of obs: 126, groups:  colony, 34
##
## Conditional model:
##                     Estimate Std. Error z value Pr(>|z|)
## (Intercept)           0.0327     0.3085   0.106    0.916
## corpse_dummyDummies   0.3130     0.3616   0.866    0.387
## dead_odourorange      0.2001     0.3618   0.553    0.580
# Pairwise drinking_time#
Pairwise3b = emmeans(Exp3bnegative_m, spec= "corpse_dummy")

Pairwise3b=contrast(Pairwise3b, method = "pairwise")
summary(Pairwise3b)
##  contrast         estimate    SE  df t.ratio p.value
##  Corpse - Dummies   -0.313 0.362 122  -0.866  0.3884
##
## Results are averaged over the levels of: dead_odour
## Results are given on the log odds ratio (not the response) scale.
# Separate treatment drinking_time#

Exp3bnegative_dummy <- subset (Exp3negative, corpse_dummy=="Dummies")
summary(Exp3bnegative_dummy)
##      colony       dead_odour         dead_side             number
##  Min.   :26.00   Length:64          Length:64          Min.   : 1.00
##  1st Qu.:38.00   Class :character   Class :character   1st Qu.: 4.75
##  Median :52.50   Mode  :character   Mode  :character   Median : 8.50
##  Mean   :54.12                                         Mean   : 8.50
##  3rd Qu.:66.75                                         3rd Qu.:12.25
##  Max.   :93.00                                         Max.   :16.00
##  corpse_dummy       first_choice       time_till_interrupt drinking_time_dead
##  Length:64          Length:64          Min.   : 4.00       Min.   : 0.00
##  Class :character   Class :character   1st Qu.:17.75       1st Qu.: 6.00
##  Mode  :character   Mode  :character   Median :25.00       Median :18.50
##                                        Mean   :28.97       Mean   :21.64
##                                        3rd Qu.:43.00       3rd Qu.:33.25
##                                        Max.   :58.00       Max.   :60.00
##  drinking_time_other  Aversiveness    Changed before 10sec    Changes
##  Min.   : 0.00       Min.   :0.0000   Min.   :0.000        Min.   : 0.000
##  1st Qu.:14.50       1st Qu.:0.0000   1st Qu.:0.000        1st Qu.: 2.000
##  Median :31.00       Median :0.0000   Median :0.000        Median : 3.000
##  Mean   :31.03       Mean   :0.4844   Mean   :0.125        Mean   : 3.438
##  3rd Qu.:47.50       3rd Qu.:1.0000   3rd Qu.:0.000        3rd Qu.: 5.000
##  Max.   :71.00       Max.   :1.0000   Max.   :1.000        Max.   :12.000
##  Drinking_aversive first_choice_feeder
##  Min.   :0.0000    Length:64
##  1st Qu.:0.0000    Class :character
##  Median :1.0000    Mode  :character
##  Mean   :0.6094
##  3rd Qu.:1.0000
##  Max.   :1.0000
Exp3bnegative_corpse <- subset (Exp3negative, corpse_dummy=="Corpse")
summary(Exp3bnegative_corpse)
##      colony       dead_odour         dead_side             number
##  Min.   :26.00   Length:62          Length:62          Min.   : 1.000
##  1st Qu.:30.00   Class :character   Class :character   1st Qu.: 5.000
##  Median :52.00   Mode  :character   Mode  :character   Median : 8.500
##  Mean   :50.05                                         Mean   : 8.452
##  3rd Qu.:63.00                                         3rd Qu.:12.000
##  Max.   :77.00                                         Max.   :16.000
##  corpse_dummy       first_choice       time_till_interrupt drinking_time_dead
##  Length:62          Length:62          Min.   : 2.00       Min.   : 0.00
##  Class :character   Class :character   1st Qu.:11.00       1st Qu.: 8.00
##  Mode  :character   Mode  :character   Median :22.00       Median :23.00
##                                        Mean   :26.15       Mean   :23.85
##                                        3rd Qu.:37.00       3rd Qu.:31.00
##                                        Max.   :78.00       Max.   :63.00
##  drinking_time_other  Aversiveness    Changed before 10sec    Changes
##  Min.   : 0.00       Min.   :0.0000   Min.   :0.0000       Min.   : 0.000
##  1st Qu.:12.75       1st Qu.:0.0000   1st Qu.:0.0000       1st Qu.: 2.250
##  Median :31.50       Median :0.0000   Median :0.0000       Median : 4.000
##  Mean   :32.69       Mean   :0.4516   Mean   :0.1774       Mean   : 4.355
##  3rd Qu.:47.75       3rd Qu.:1.0000   3rd Qu.:0.0000       3rd Qu.: 5.000
##  Max.   :93.00       Max.   :1.0000   Max.   :1.0000       Max.   :13.000
##  Drinking_aversive first_choice_feeder
##  Min.   :0.0000    Length:62
##  1st Qu.:0.0000    Class :character
##  Median :1.0000    Mode  :character
##  Mean   :0.5323
##  3rd Qu.:1.0000
##  Max.   :1.0000
Exp3bnegative_dummy_m <-glmer(Drinking_aversive ~ 1 + (1|colony),
                                             family="binomial",
                                             data= Exp3bnegative_dummy )
## boundary (singular) fit: see help('isSingular')
summary(Exp3bnegative_dummy_m)
## Generalized linear mixed model fit by maximum likelihood (Laplace
##   Approximation) [glmerMod]
##  Family: binomial  ( logit )
## Formula: Drinking_aversive ~ 1 + (1 | colony)
##    Data: Exp3bnegative_dummy
##
##      AIC      BIC   logLik deviance df.resid
##     89.6     94.0    -42.8     85.6       62
##
## Scaled residuals:
##     Min      1Q  Median      3Q     Max
## -1.2490 -1.2490  0.8006  0.8006  0.8006
##
## Random effects:
##  Groups Name        Variance Std.Dev.
##  colony (Intercept) 0        0
## Number of obs: 64, groups:  colony, 19
##
## Fixed effects:
##             Estimate Std. Error z value Pr(>|z|)
## (Intercept)   0.4447     0.2562   1.736   0.0826 .
## ---
## Signif. codes:  0 '***' 0.001 '**' 0.01 '*' 0.05 '.' 0.1 ' ' 1
## optimizer (Nelder_Mead) convergence code: 0 (OK)
## boundary (singular) fit: see help('isSingular')
Exp3bnegative_corpse_m <-glmer(Drinking_aversive ~ 1 + (1|colony),
                                             family="binomial",
                                             data= Exp3bnegative_corpse )
## boundary (singular) fit: see help('isSingular')
summary(Exp3bnegative_corpse_m)
## Generalized linear mixed model fit by maximum likelihood (Laplace
##   Approximation) [glmerMod]
##  Family: binomial  ( logit )
## Formula: Drinking_aversive ~ 1 + (1 | colony)
##    Data: Exp3bnegative_corpse
##
##      AIC      BIC   logLik deviance df.resid
##     89.7     93.9    -42.8     85.7       60
##
## Scaled residuals:
##     Min      1Q  Median      3Q     Max
## -1.0667 -1.0667  0.9374  0.9374  0.9374
##
## Random effects:
##  Groups Name        Variance Std.Dev.
##  colony (Intercept) 0        0
## Number of obs: 62, groups:  colony, 18
##
## Fixed effects:
##             Estimate Std. Error z value Pr(>|z|)
## (Intercept)   0.1292     0.2545   0.508    0.612
## optimizer (Nelder_Mead) convergence code: 0 (OK)
## boundary (singular) fit: see help('isSingular')
# 4 days starvation


# GLMM drinking_time

Exp3_drinking_negative4days_m <-glmmTMB(Drinking_aversive ~ corpse_dummy + dead_odour + (1|Colony_ID),
                              family="binomial",
                              data= Exp3negative4days)
summary(Exp3_drinking_negative4days_m)
##  Family: binomial  ( logit )
## Formula:
## Drinking_aversive ~ corpse_dummy + dead_odour + (1 | Colony_ID)
## Data: Exp3negative4days
##
##      AIC      BIC   logLik deviance df.resid
##    190.0    201.6    -91.0    182.0      129
##
## Random effects:
##
## Conditional model:
##  Groups    Name        Variance Std.Dev.
##  Colony_ID (Intercept) 0.05817  0.2412
## Number of obs: 133, groups:  Colony_ID, 5
##
## Conditional model:
##                      Estimate Std. Error z value Pr(>|z|)
## (Intercept)           -0.2988     0.3641  -0.821    0.412
## corpse_dummydummy      0.5325     0.3560   1.496    0.135
## dead_odourstrawberry   0.1087     0.4345   0.250    0.802
# Separate treatment for drinking time

Exp3b_drinking_negative4days_dummy<- subset (Exp3negative4days, corpse_dummy=="dummy")

Exp3b_drinking_negative4days_corpse <- subset (Exp3negative4days, corpse_dummy=="corpse")


Exp3b_drinking_negative4days_dummy_m <-glmer(Drinking_aversive ~ 1 + (1|Colony_ID),
                                             family="binomial",
                                             data= Exp3b_drinking_negative4days_dummy)
summary(Exp3b_drinking_negative4days_dummy_m)
## Generalized linear mixed model fit by maximum likelihood (Laplace
##   Approximation) [glmerMod]
##  Family: binomial  ( logit )
## Formula: Drinking_aversive ~ 1 + (1 | Colony_ID)
##    Data: Exp3b_drinking_negative4days_dummy
##
##      AIC      BIC   logLik deviance df.resid
##     95.1     99.5    -45.5     91.1       66
##
## Scaled residuals:
##     Min      1Q  Median      3Q     Max
## -1.4198 -1.0430  0.7043  0.7391  1.4331
##
## Random effects:
##  Groups    Name        Variance Std.Dev.
##  Colony_ID (Intercept) 0.4863   0.6974
## Number of obs: 68, groups:  Colony_ID, 5
##
## Fixed effects:
##             Estimate Std. Error z value Pr(>|z|)
## (Intercept)   0.2274     0.4097   0.555    0.579
Exp3b_drinking_negative4days_corpse_m <-glmer(Drinking_aversive ~ 1 + (1|Colony_ID),
                                             family="binomial",
                                             data= Exp3b_drinking_negative4days_corpse)
## boundary (singular) fit: see help('isSingular')
summary(Exp3b_drinking_negative4days_corpse_m)
## Generalized linear mixed model fit by maximum likelihood (Laplace
##   Approximation) [glmerMod]
##  Family: binomial  ( logit )
## Formula: Drinking_aversive ~ 1 + (1 | Colony_ID)
##    Data: Exp3b_drinking_negative4days_corpse
##
##      AIC      BIC   logLik deviance df.resid
##     93.4     97.7    -44.7     89.4       63
##
## Scaled residuals:
##     Min      1Q  Median      3Q     Max
## -0.8975 -0.8975 -0.8975  1.1142  1.1142
##
## Random effects:
##  Groups    Name        Variance Std.Dev.
##  Colony_ID (Intercept) 0        0
## Number of obs: 65, groups:  Colony_ID, 5
##
## Fixed effects:
##             Estimate Std. Error z value Pr(>|z|)
## (Intercept)  -0.2162     0.2495  -0.867    0.386
## optimizer (Nelder_Mead) convergence code: 0 (OK)
## boundary (singular) fit: see help('isSingular')
summary(Exp3b_drinking_negative4days_corpse)
##   dead_odour         dead_side             Ant_id      corpse_dummy
##  Length:65          Length:65          Min.   :  1.0   Length:65
##  Class :character   Class :character   1st Qu.: 26.0   Class :character
##  Mode  :character   Mode  :character   Median : 56.0   Mode  :character
##                                        Mean   : 60.8
##                                        3rd Qu.: 87.0
##                                        Max.   :118.0
##  first_choice       time_till_first_interrupt drinking_time_dead
##  Length:65          Min.   :  0.50            Min.   :  0.00
##  Class :character   1st Qu.: 35.00            1st Qu.:  9.00
##  Mode  :character   Median : 54.00            Median : 39.00
##                     Mean   : 54.79            Mean   : 42.94
##                     3rd Qu.: 76.00            3rd Qu.: 68.00
##                     Max.   :114.00            Max.   :133.00
##  drinking_time_other  Aversiveness    Changed before 10sec    Changes
##  Min.   :  0.00      Min.   :0.0000   Min.   :0.00000      Min.   : 0.000
##  1st Qu.: 12.00      1st Qu.:0.0000   1st Qu.:0.00000      1st Qu.: 2.000
##  Median : 30.00      Median :0.0000   Median :0.00000      Median : 3.000
##  Mean   : 42.22      Mean   :0.4923   Mean   :0.07692      Mean   : 4.246
##  3rd Qu.: 73.00      3rd Qu.:1.0000   3rd Qu.:0.00000      3rd Qu.: 6.000
##  Max.   :144.00      Max.   :1.0000   Max.   :1.00000      Max.   :19.000
##    Colony_ID     Drinking_aversive
##  Min.   :41.00   Min.   :0.0000
##  1st Qu.:44.00   1st Qu.:0.0000
##  Median :64.00   Median :0.0000
##  Mean   :54.68   Mean   :0.4462
##  3rd Qu.:65.00   3rd Qu.:1.0000
##  Max.   :66.00   Max.   :1.0000
```

- Ants did not drink significantly shorter on the feeder flavoured with the corpse odour.
- There is no difference between the two treatments dummies and corpses.

Does this model work?

dharmit3b <-simulateResiduals(Exp3bnegative_m)
plot(dharmit3b)

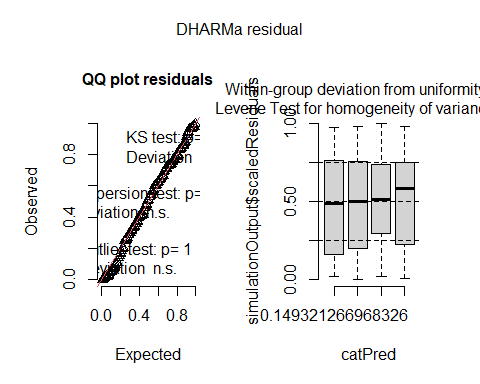
 - Yes it works.

##### Figures

```
## drinking_time, 2 days starvation

fig3cN <- ggplot (Exp3negative, aes(x = corpse_dummy, y = Drinking_aversive ))+
  scale_y_continuous(expand = c(0, 0)) + # forces X axis to 0, but in this case is overriden by ribbon
  geom_point( alpha = 0) +
  stat_summary(fun.y = "mean", geom = "bar", fill="dodgerblue4") +
  ylab("Proportion of choices for non-corpse odour") +
  xlab("Treatment") + stat_summary(fun.data = "mean_cl_boot", geom="errorbar", width = 0.2) +
  theme_bw(18) +
  geom_abline(intercept = 0.5, slope = 0, color = "black", linetype = 2) +
  coord_cartesian( ylim = c(0, 1.0)) # zooms in on the top half
## Warning: `fun.y` is deprecated. Use `fun` instead.
print(fig3cN)
```

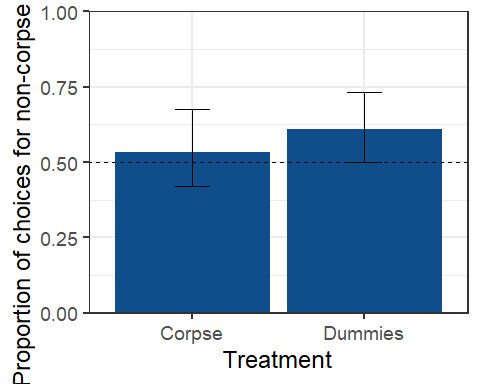

```
## drinking_time, 4 days starvation

fig3dN <- ggplot (Exp3negative4days, aes(x = corpse_dummy, y = Drinking_aversive ))+
  scale_y_continuous(expand = c(0, 0)) + # forces X axis to 0, but in this case is overriden by ribbon
  geom_point( alpha = 0) +
  stat_summary(fun.y = "mean", geom = "bar", fill="dodgerblue4") +
  ylab("Proportion of choices for non-corpse odour") +
  xlab("Treatment") + stat_summary(fun.data = "mean_cl_boot", geom="errorbar", width = 0.2) +
  theme_bw(18) +
  geom_abline(intercept = 0.5, slope = 0, color = "black", linetype = 2) +
  coord_cartesian( ylim = c(0, 1.0)) # zooms in on the top half
## Warning: `fun.y` is deprecated. Use `fun` instead.
print(fig3dN)
```

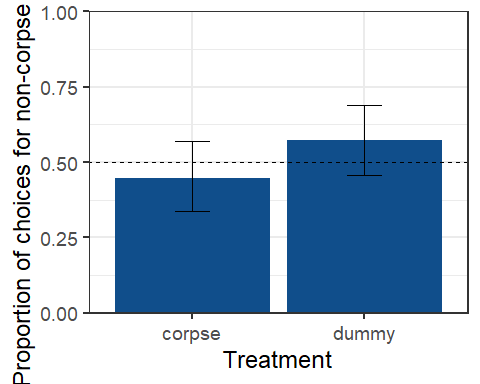

```
summary(Exp3negative4days)
##   dead_odour         dead_side             Ant_id    corpse_dummy
##  Length:133         Length:133         Min.   :  1   Length:133
##  Class :character   Class :character   1st Qu.: 34   Class :character
##  Mode  :character   Mode  :character   Median : 67   Mode  :character
##                                        Mean   : 67
##                                        3rd Qu.:100
##                                        Max.   :133
##  first_choice       time_till_first_interrupt drinking_time_dead
##  Length:133         Min.   :  0.50            Min.   :  0.00
##  Class :character   1st Qu.: 34.00            1st Qu.:  5.00
##  Mode  :character   Median : 48.00            Median : 32.00
##                     Mean   : 51.44            Mean   : 37.05
##                     3rd Qu.: 69.00            3rd Qu.: 63.00
##                     Max.   :124.00            Max.   :133.00
##  drinking_time_other  Aversiveness    Changed before 10sec    Changes
##  Min.   :  0.00      Min.   :0.0000   Min.   :0.00000      Min.   : 0.000
##  1st Qu.: 10.00      1st Qu.:0.0000   1st Qu.:0.00000      1st Qu.: 2.000
##  Median : 33.00      Median :1.0000   Median :0.00000      Median : 3.000
##  Mean   : 43.91      Mean   :0.5414   Mean   :0.05263      Mean   : 4.143
##  3rd Qu.: 71.00      3rd Qu.:1.0000   3rd Qu.:0.00000      3rd Qu.: 6.000
##  Max.   :149.00      Max.   :1.0000   Max.   :1.00000      Max.   :19.000
##    Colony_ID     Drinking_aversive
##  Min.   :41.00   Min.   :0.0000
##  1st Qu.:44.00   1st Qu.:0.0000
##  Median :64.00   Median :1.0000
##  Mean   :54.87   Mean   :0.5113
##  3rd Qu.:65.00   3rd Qu.:1.0000
##  Max.   :66.00   Max.   :1.0000
summary(Exp3negative)
##      colony       dead_odour         dead_side             number
##  Min.   :26.00   Length:126         Length:126         Min.   : 1.000
##  1st Qu.:30.00   Class :character   Class :character   1st Qu.: 5.000
##  Median :52.00   Mode  :character   Mode  :character   Median : 8.500
##  Mean   :52.12                                         Mean   : 8.476
##  3rd Qu.:63.00                                         3rd Qu.:12.000
##  Max.   :93.00                                         Max.   :16.000
##  corpse_dummy       first_choice       time_till_interrupt drinking_time_dead
##  Length:126         Length:126         Min.   : 2.00       Min.   : 0.00
##  Class :character   Class :character   1st Qu.:14.00       1st Qu.: 7.25
##  Mode  :character   Mode  :character   Median :24.00       Median :22.00
##                                        Mean   :27.58       Mean   :22.73
##                                        3rd Qu.:38.75       3rd Qu.:33.00
##                                        Max.   :78.00       Max.   :63.00
##  drinking_time_other  Aversiveness    Changed before 10sec    Changes
##  Min.   : 0.00       Min.   :0.0000   Min.   :0.0000       Min.   : 0.000
##  1st Qu.:13.50       1st Qu.:0.0000   1st Qu.:0.0000       1st Qu.: 2.000
##  Median :31.00       Median :0.0000   Median :0.0000       Median : 3.000
##  Mean   :31.85       Mean   :0.4683   Mean   :0.1508       Mean   : 3.889
##  3rd Qu.:47.75       3rd Qu.:1.0000   3rd Qu.:0.0000       3rd Qu.: 5.000
##  Max.   :93.00       Max.   :1.0000   Max.   :1.0000       Max.   :13.000
##  Drinking_aversive first_choice_feeder
##  Min.   :0.0000    Length:126
##  1st Qu.:0.0000    Class :character
##  Median :1.0000    Mode  :character
##  Mean   :0.5714
##  3rd Qu.:1.0000
##  Max.   :1.0000
```

##### Merged Figures exp 1-2 and all exp 3 figures

```
## Plot exp 1-2 experiments into 2 panels ##


fig2N <- fig2N + ylab("")
Individual12plot <- grid.arrange(fig1N, fig2N, nrow = 1)
```

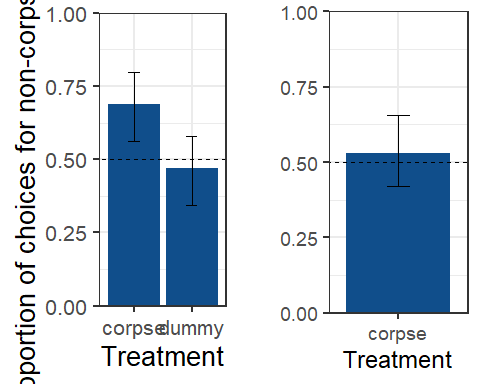

```
print(Individual12plot)
## TableGrob (1 x 2) "arrange": 2 grobs
##   z     cells    name           grob
## 1 1 (1-1,1-1) arrange gtable[layout]
## 2 2 (1-1,2-2) arrange gtable[layout]
##for powerpoint export##

ggsave ("Individual12plot.png", plot = Individual12plot, dpi = 300, width = 20, height = 20, units = c("cm"))

read_pptx() %>%
add_slide(layout = "Title and Content", master = "Office Theme") %>%
  ph_with(value = dml(grid.arrange(fig1N, fig2N,nrow = 1)), location = ph_location_type(type = "body")) %>%
 print(target = "Individual12.pptx") %>%
  browseURL()

## Plot exp 3, 2 and 4 days starvation into 2 panels#


fig3dN <- fig3dN + ylab("")
Individual3plot <- grid.arrange(fig3cN, fig3dN, nrow = 1)
```

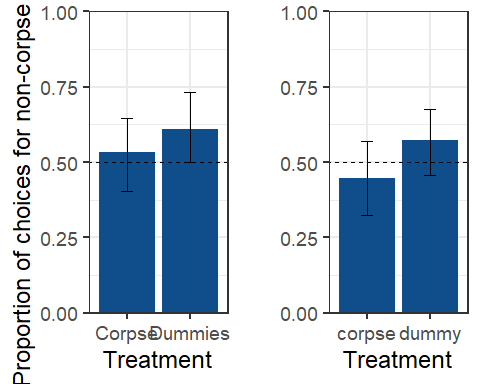

```
##for powerpoint export##

ggsave ("Individual3.png", plot = Individual3plot, dpi = 300, width = 20, height = 20, units = c("cm"))

read_pptx() %>%
add_slide(layout = "Title and Content", master = "Office Theme") %>%
  ph_with(value = dml(grid.arrange(fig3cN,fig3dN, nrow = 1)), location = ph_location_type(type = "body")) %>%
 print(target = "Individual3.pptx") %>%
  browseURL()
```

#### 4. Analysis, Experiment 4 – Scented corpses, bifurcated choice flavoured food hydrogel beads

##### Glmm modeling, pairwise, Experiment 4, Can the presence of scented corpses drive collective food choice?

```
# For supplement, with all colonies

Exp4Recovery <- read_excel("C:/Users/LocalAdmin/Documents/ThomasWagnerPhD/PhD/RawData/OvercomingNegativStimuli/All4Exp/Exp4Collective/Exp4Recovery.xlsx")


Exp4Recovery $feedertype<-as.factor(Exp4Recovery$feedertype)
Exp4Recovery $antsOnFeeder<-as.numeric(Exp4Recovery$antsOnFeeder)

## subdata#
Exp4RecoveryOnly60m <-subset(Exp4Recovery, Time_after_start_min < 120)
Exp4Recovery23h <-subset(Exp4Recovery, Time_after_start_min > 119)


#Glmm Ants staying/feeding on the two different feeders within 60min


Exp4RecoveryOnly60m_m <-glmmTMB(antsOnFeeder ~ feedertype + Time_after_start_min + Dead_odour + (1|Colony), ziformula = ~1,
                              family= poisson,
                              data= Exp4RecoveryOnly60m)

summary(Exp4RecoveryOnly60m_m)
##  Family: poisson  ( log )
## Formula:
## antsOnFeeder ~ feedertype + Time_after_start_min + Dead_odour +
##     (1 | Colony)
## Zero inflation:                ~1
## Data: Exp4RecoveryOnly60m
##
##      AIC      BIC   logLik deviance df.resid
##   2595.5   2619.5  -1291.8   2583.5      396
##
## Random effects:
##
## Conditional model:
##  Groups Name        Variance Std.Dev.
##  Colony (Intercept) 1.145    1.07
## Number of obs: 402, groups:  Colony, 16
##
## Conditional model:
##                       Estimate Std. Error z value Pr(>|z|)
## (Intercept)           1.349407   0.387469   3.483 0.000497 ***
## feedertypenovel       0.385358   0.051792   7.440 1.00e-13 ***
## Time_after_start_min -0.007393   0.001367  -5.407 6.41e-08 ***
## Dead_odourStrawberry  0.090973   0.542846   0.168 0.866910
## ---
## Signif. codes:  0 '***' 0.001 '**' 0.01 '*' 0.05 '.' 0.1 ' ' 1
##
## Zero-inflation model:
##             Estimate Std. Error z value Pr(>|z|)
## (Intercept)  -1.1026     0.1361  -8.102 5.41e-16 ***
## ---
## Signif. codes:  0 '***' 0.001 '**' 0.01 '*' 0.05 '.' 0.1 ' ' 1
dharmitRecover <-simulateResiduals(Exp4RecoveryOnly60m_m)
plot(dharmitRecover)
```

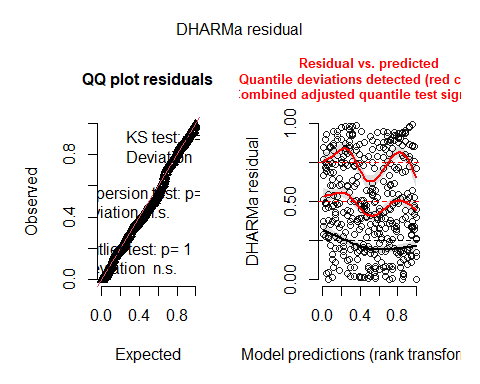

```
#Pairwise 60min

PairwiseAllColoniesAntsFeeder60mBar = emmeans(Exp4RecoveryOnly60m_m, spec= "feedertype")

PairwiseAllColoniesAntsFeeder60mBar=contrast(PairwiseAllColoniesAntsFeeder60mBar, method = "pairwise")
summary(PairwiseAllColoniesAntsFeeder60mBar)
##  contrast     estimate     SE  df t.ratio p.value
##  dead - novel   -0.385 0.0518 396  -7.440  <.0001
##
## Results are averaged over the levels of: Dead_odour
## Results are given on the log (not the response) scale.
#Glmm Ants staying/feeding on the two different feeders over 23h

Exp4Recovery23h_m <-glmmTMB(antsOnFeeder ~ feedertype + Time_after_start_min + Dead_odour + (1|Colony), ziformula = ~1,
                              family= poisson,
                              data= Exp4Recovery23h)
## Warning in (function (start, objective, gradient = NULL, hessian = NULL, : NA/
## NaN function evaluation
summary(Exp4Recovery23h_m)
##  Family: poisson  ( log )
## Formula:
## antsOnFeeder ~ feedertype + Time_after_start_min + Dead_odour +
##     (1 | Colony)
## Zero inflation:                ~1
## Data: Exp4Recovery23h
##
##      AIC      BIC   logLik deviance df.resid
##    592.4    618.7   -290.2    580.4      584
##
## Random effects:
##
## Conditional model:
##  Groups Name        Variance Std.Dev.
##  Colony (Intercept) 0.5466   0.7393
## Number of obs: 590, groups:  Colony, 16
##
## Conditional model:
##                        Estimate Std. Error z value Pr(>|z|)
## (Intercept)          -0.2891742  0.3850426  -0.751    0.453
## feedertypenovel       0.3366789  0.2168297   1.553    0.120
## Time_after_start_min -0.0015622  0.0002703  -5.779 7.52e-09 ***
## Dead_odourStrawberry  0.5460937  0.4413972   1.237    0.216
## ---
## Signif. codes:  0 '***' 0.001 '**' 0.01 '*' 0.05 '.' 0.1 ' ' 1
##
## Zero-inflation model:
##             Estimate Std. Error z value Pr(>|z|)
## (Intercept)   0.2525     0.2757   0.915     0.36
dharmitRecover2 <-simulateResiduals(Exp4Recovery23h_m)
plot(dharmitRecover2)
```

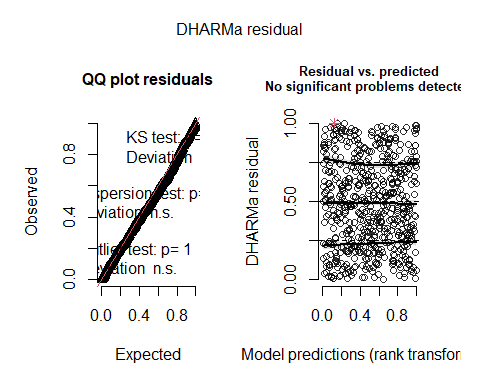

```
#pairwise 23h
PairwiseAllColoniesAntsFeeder23hBar = emmeans(Exp4Recovery23h_m, spec= "feedertype")

PairwiseAllColoniesAntsFeeder23hBar=contrast(PairwiseAllColoniesAntsFeeder23hBar, method = "pairwise")
summary(PairwiseAllColoniesAntsFeeder23hBar)
##  contrast     estimate    SE  df t.ratio p.value
##  dead - novel   -0.337 0.217 584  -1.553  0.1210
##
## Results are averaged over the levels of: Dead_odour
## Results are given on the log (not the response) scale.
# Proportion, subdata #

Exp4Prob60m<- subset(Exp4Prob, Time_after_start_min < 100)
Exp4Prob60m$feedertype <-as.factor(Exp4Prob60m$feedertype)
Exp4Prob60m$prop.on.feeder <-as.numeric(Exp4Prob60m$prop.on.feeder)
Exp4Prob60m$Colony <- as.factor(Exp4Prob60m$Colony)
Exp4Prob60m <- na.omit(Exp4Prob60m)
Exp4Prob60m$Colony <- as.numeric(gsub("[^0-9]", "", Exp4Prob60m$Colony))

Exp4Prob23h<- subset(Exp4Prob, Time_after_start_min > 100)
Exp4Prob23h$feedertype <-as.factor(Exp4Prob23h$feedertype)
Exp4Prob23h$prop.on.feeder <-as.numeric(Exp4Prob23h$prop.on.feeder)
Exp4Prob23h$Colony <- as.factor(Exp4Prob23h$Colony)
Exp4Prob23h<- na.omit(Exp4Prob23h)
Exp4Prob23h$Colony <- as.numeric(gsub("[^0-9]", "", Exp4Prob23h$Colony))

# GLMM Proportion of ants feeding over 60m on a specific feeder


# Fit the zero-inflated beta regression model
zero_inflated_model_60m <- gamlss(prop.on.feeder ~ feedertype + Time_after_start_min + Dead_odour,
                              family = BEINF,
                              data = Exp4Prob60m)
## GAMLSS-RS iteration 1: Global Deviance = 725.5363
## GAMLSS-RS iteration 2: Global Deviance = 724.1927
## GAMLSS-RS iteration 3: Global Deviance = 724.1775
## GAMLSS-RS iteration 4: Global Deviance = 724.1773
# Print the model summary
summary(zero_inflated_model_60m)
## ******************************************************************
## Family:  c("BEINF", "Beta Inflated")
##
## Call:  gamlss(formula = prop.on.feeder ~ feedertype + Time_after_start_min +
##     Dead_odour, family = BEINF, data = Exp4Prob60m)
##
## Fitting method: RS()
##
## ------------------------------------------------------------------
## Mu link function:  logit
## Mu Coefficients:
##                        Estimate Std. Error t value Pr(>|t|)
## (Intercept)          -2.203e-01  2.081e-01  -1.058  0.29066
## feedertypenovel       4.405e-01  1.586e-01   2.777  0.00578 **
## Time_after_start_min -2.437e-18  4.622e-03   0.000  1.00000
## Dead_odourStrawberry -2.164e-16  1.612e-01   0.000  1.00000
## ---
## Signif. codes:  0 '***' 0.001 '**' 0.01 '*' 0.05 '.' 0.1 ' ' 1
##
## ------------------------------------------------------------------
## Sigma link function:  logit
## Sigma Coefficients:
##             Estimate Std. Error t value Pr(>|t|)
## (Intercept)  0.15258    0.07098    2.15   0.0323 *
## ---
## Signif. codes:  0 '***' 0.001 '**' 0.01 '*' 0.05 '.' 0.1 ' ' 1
##
## ------------------------------------------------------------------
## Nu link function:  log
## Nu Coefficients:
##             Estimate Std. Error t value Pr(>|t|)
## (Intercept)  -0.6706     0.1303  -5.146 4.48e-07 ***
## ---
## Signif. codes:  0 '***' 0.001 '**' 0.01 '*' 0.05 '.' 0.1 ' ' 1
##
## ------------------------------------------------------------------
## Tau link function:  log
## Tau Coefficients:
##             Estimate Std. Error t value Pr(>|t|)
## (Intercept)  -0.6705     0.1303  -5.145  4.5e-07 ***
## ---
## Signif. codes:  0 '***' 0.001 '**' 0.01 '*' 0.05 '.' 0.1 ' ' 1
##
## ------------------------------------------------------------------
## No. of observations in the fit:  352
## Degrees of Freedom for the fit:  7
##       Residual Deg. of Freedom:  345
##                       at cycle:  4
##
## Global Deviance:     724.1773
##             AIC:     738.1773
##             SBC:     765.2227
## ******************************************************************
# Pairwise
PairwiseAllColoniesAntsFeeder60m = emmeans(zero_inflated_model_60m, spec= "feedertype")
## ******************************************************************
## Family:  c("BEINF", "Beta Inflated")
##
## Call:  gamlss(formula = prop.on.feeder ~ feedertype + Time_after_start_min +
##     Dead_odour, family = BEINF, data = Exp4Prob60m)
##
## Fitting method: RS()
##
## ------------------------------------------------------------------
## Mu link function:  logit
## Mu Coefficients:
##                        Estimate Std. Error t value Pr(>|t|)
## (Intercept)          -2.203e-01  2.081e-01  -1.058  0.29066
## feedertypenovel       4.405e-01  1.586e-01   2.777  0.00578 **
## Time_after_start_min -2.437e-18  4.622e-03   0.000  1.00000
## Dead_odourStrawberry -2.164e-16  1.612e-01   0.000  1.00000
## ---
## Signif. codes:  0 '***' 0.001 '**' 0.01 '*' 0.05 '.' 0.1 ' ' 1
##
## ------------------------------------------------------------------
## Sigma link function:  logit
## Sigma Coefficients:
##             Estimate Std. Error t value Pr(>|t|)
## (Intercept)  0.15258    0.07098    2.15   0.0323 *
## ---
## Signif. codes:  0 '***' 0.001 '**' 0.01 '*' 0.05 '.' 0.1 ' ' 1
##
## ------------------------------------------------------------------
## Nu link function:  log
## Nu Coefficients:
##             Estimate Std. Error t value Pr(>|t|)
## (Intercept)  -0.6706     0.1303  -5.146 4.48e-07 ***
## ---
## Signif. codes:  0 '***' 0.001 '**' 0.01 '*' 0.05 '.' 0.1 ' ' 1
##
## ------------------------------------------------------------------
## Tau link function:  log
## Tau Coefficients:
##             Estimate Std. Error t value Pr(>|t|)
## (Intercept)  -0.6705     0.1303  -5.145  4.5e-07 ***
## ---
## Signif. codes:  0 '***' 0.001 '**' 0.01 '*' 0.05 '.' 0.1 ' ' 1
##
## ------------------------------------------------------------------
## No. of observations in the fit:  352
## Degrees of Freedom for the fit:  7
##       Residual Deg. of Freedom:  345
##                       at cycle:  4
##
## Global Deviance:     724.1773
##             AIC:     738.1773
##             SBC:     765.2227
## ******************************************************************
PairwiseAllColoniesAntsFeeder60m=contrast(PairwiseAllColoniesAntsFeeder60m, method = "pairwise")
summary(PairwiseAllColoniesAntsFeeder60m)
##  contrast     estimate    SE  df z.ratio p.value
##  dead - novel   -0.441 0.159 Inf  -2.777  0.0055
##
## Results are averaged over the levels of: Dead_odour
## Results are given on the log odds ratio (not the response) scale.
# GLMM Proportion of ants feeding over 23h on a specific feeder

# Fit the zero-inflated beta regression model
zero_inflated_model_23h_m <- gamlss(prop.on.feeder ~ feedertype + Time_after_start_min + Dead_odour,
                              family = BEINF,
                              data = Exp4Prob23h)
## GAMLSS-RS iteration 1: Global Deviance = 290.2294
## GAMLSS-RS iteration 2: Global Deviance = 264.3923
## GAMLSS-RS iteration 3: Global Deviance = 259.5146
## GAMLSS-RS iteration 4: Global Deviance = 258.0431
## GAMLSS-RS iteration 5: Global Deviance = 257.5125
## GAMLSS-RS iteration 6: Global Deviance = 257.3029
## GAMLSS-RS iteration 7: Global Deviance = 257.2158
## GAMLSS-RS iteration 8: Global Deviance = 257.1783
## GAMLSS-RS iteration 9: Global Deviance = 257.1619
## GAMLSS-RS iteration 10: Global Deviance = 257.1547
## GAMLSS-RS iteration 11: Global Deviance = 257.1514
## GAMLSS-RS iteration 12: Global Deviance = 257.1499
## GAMLSS-RS iteration 13: Global Deviance = 257.1493
# Print the model summary
summary(zero_inflated_model_23h_m)
## ******************************************************************
## Family:  c("BEINF", "Beta Inflated")
##
## Call:  gamlss(formula = prop.on.feeder ~ feedertype + Time_after_start_min +
##     Dead_odour, family = BEINF, data = Exp4Prob23h)
##
## Fitting method: RS()
##
## ------------------------------------------------------------------
## Mu link function:  logit
## Mu Coefficients:
##                        Estimate Std. Error t value Pr(>|t|)
## (Intercept)          -2.496e-01  3.141e-01  -0.795    0.428
## feedertypenovel       4.992e-01  3.312e-01   1.508    0.134
## Time_after_start_min -5.632e-18  5.363e-04   0.000    1.000
## Dead_odourStrawberry  4.709e-15  3.742e-01   0.000    1.000
##
## ------------------------------------------------------------------
## Sigma link function:  logit
## Sigma Coefficients:
##             Estimate Std. Error t value Pr(>|t|)
## (Intercept)  -0.8077     0.2355  -3.429 0.000801 ***
## ---
## Signif. codes:  0 '***' 0.001 '**' 0.01 '*' 0.05 '.' 0.1 ' ' 1
##
## ------------------------------------------------------------------
## Nu link function:  log
## Nu Coefficients:
##             Estimate Std. Error t value Pr(>|t|)
## (Intercept)   1.4811     0.2953   5.016 1.62e-06 ***
## ---
## Signif. codes:  0 '***' 0.001 '**' 0.01 '*' 0.05 '.' 0.1 ' ' 1
##
## ------------------------------------------------------------------
## Tau link function:  log
## Tau Coefficients:
##             Estimate Std. Error t value Pr(>|t|)
## (Intercept)   1.5600     0.2932   5.321 4.14e-07 ***
## ---
## Signif. codes:  0 '***' 0.001 '**' 0.01 '*' 0.05 '.' 0.1 ' ' 1
##
## ------------------------------------------------------------------
## No. of observations in the fit:  143
## Degrees of Freedom for the fit:  7
##       Residual Deg. of Freedom:  136
##                       at cycle:  13
##
## Global Deviance:     257.1493
##             AIC:     271.1493
##             SBC:     291.8892
## ******************************************************************
# Pairwise
PairwiseAllColoniesAntsFeeder23h = emmeans(zero_inflated_model_23h_m, spec= "feedertype")
## ******************************************************************
## Family:  c("BEINF", "Beta Inflated")
##
## Call:  gamlss(formula = prop.on.feeder ~ feedertype + Time_after_start_min +
##     Dead_odour, family = BEINF, data = Exp4Prob23h)
##
## Fitting method: RS()
##
## ------------------------------------------------------------------
## Mu link function:  logit
## Mu Coefficients:
##                        Estimate Std. Error t value Pr(>|t|)
## (Intercept)          -2.496e-01  3.141e-01  -0.795    0.428
## feedertypenovel       4.992e-01  3.312e-01   1.508    0.134
## Time_after_start_min -5.632e-18  5.363e-04   0.000    1.000
## Dead_odourStrawberry  4.709e-15  3.742e-01   0.000    1.000
##
## ------------------------------------------------------------------
## Sigma link function:  logit
## Sigma Coefficients:
##             Estimate Std. Error t value Pr(>|t|)
## (Intercept)  -0.8077     0.2355  -3.429 0.000801 ***
## ---
## Signif. codes:  0 '***' 0.001 '**' 0.01 '*' 0.05 '.' 0.1 ' ' 1
##
## ------------------------------------------------------------------
## Nu link function:  log
## Nu Coefficients:
##             Estimate Std. Error t value Pr(>|t|)
## (Intercept)   1.4811     0.2953   5.016 1.62e-06 ***
## ---
## Signif. codes:  0 '***' 0.001 '**' 0.01 '*' 0.05 '.' 0.1 ' ' 1
##
## ------------------------------------------------------------------
## Tau link function:  log
## Tau Coefficients:
##             Estimate Std. Error t value Pr(>|t|)
## (Intercept)   1.5600     0.2932   5.321 4.14e-07 ***
## ---
## Signif. codes:  0 '***' 0.001 '**' 0.01 '*' 0.05 '.' 0.1 ' ' 1
##
## ------------------------------------------------------------------
## No. of observations in the fit:  143
## Degrees of Freedom for the fit:  7
##       Residual Deg. of Freedom:  136
##                       at cycle:  13
##
## Global Deviance:     257.1493
##             AIC:     271.1493
##             SBC:     291.8892
## ******************************************************************
PairwiseAllColoniesAntsFeeder23h=contrast(PairwiseAllColoniesAntsFeeder23h, method = "pairwise")
summary(PairwiseAllColoniesAntsFeeder23h)
##  contrast     estimate    SE  df z.ratio p.value
##  dead - novel   -0.499 0.331 Inf  -1.508  0.1317
##
## Results are averaged over the levels of: Dead_odour
## Results are given on the log odds ratio (not the response) scale.
##############################################################

# For manuscript, without too small colonies


# Read the data from Excel file
Exp4WithoutSmall <- read_excel("C:/Users/LocalAdmin/Documents/ThomasWagnerPhD/PhD/RawData/OvercomingNegativStimuli/All4Exp/Exp4Collective/Exp4Recovery.xlsx")

Exp4WithoutSmall$feedertype <- as.factor(Exp4WithoutSmall$feedertype)
Exp4WithoutSmall$Colony <- as.factor(Exp4WithoutSmall$Colony)

Exp4WithoutSmall$antsOnFeeder <- as.numeric(Exp4WithoutSmall$antsOnFeeder)
Exp4WithoutSmall$prop.on.feeder <- as.numeric(Exp4WithoutSmall$prop.on.feeder)
## Warning: NAs durch Umwandlung erzeugt
Exp4WithoutSmall <- subset(Exp4WithoutSmall, !is.infinite(prop.on.feeder))

Exp4WithoutSmall$Colony <- as.numeric(gsub("[^0-9]", "", Exp4WithoutSmall$Colony))


## subdata

Exp460mWithoutSmall <-subset(Exp4WithoutSmall, Time_after_start_min < 120 & total.feeding.ants.whole.exp > 40)

Exp423hWithoutSmall <-subset(Exp4WithoutSmall, Time_after_start_min > 119 & total.feeding.ants.whole.exp > 40)


#Glmm Ants staying/feeding on the two different feeders within 60min

Exp460mWithoutSmall_m <-glmmTMB(antsOnFeeder ~ feedertype + Time_after_start_min + Dead_odour + (1|Colony), ziformula = ~1,
                              family= poisson,
                              data= Exp460mWithoutSmall)

summary(Exp460mWithoutSmall_m)
##  Family: poisson  ( log )
## Formula:
## antsOnFeeder ~ feedertype + Time_after_start_min + Dead_odour +
##     (1 | Colony)
## Zero inflation:                ~1
## Data: Exp460mWithoutSmall
##
##      AIC      BIC   logLik deviance df.resid
##   2256.3   2277.9  -1122.1   2244.3      266
##
## Random effects:
##
## Conditional model:
##  Groups Name        Variance Std.Dev.
##  Colony (Intercept) 0.2776   0.5268
## Number of obs: 272, groups:  Colony, 8
##
## Conditional model:
##                       Estimate Std. Error z value Pr(>|z|)
## (Intercept)           1.772135   0.199184   8.897  < 2e-16 ***
## feedertypenovel       0.403350   0.052037   7.751 9.10e-15 ***
## Time_after_start_min -0.007108   0.001403  -5.065 4.07e-07 ***
## Dead_odourStrawberry  0.271996   0.064184   4.238 2.26e-05 ***
## ---
## Signif. codes:  0 '***' 0.001 '**' 0.01 '*' 0.05 '.' 0.1 ' ' 1
##
## Zero-inflation model:
##             Estimate Std. Error z value Pr(>|z|)
## (Intercept)   -1.301      0.155  -8.393   <2e-16 ***
## ---
## Signif. codes:  0 '***' 0.001 '**' 0.01 '*' 0.05 '.' 0.1 ' ' 1
# Pairwise
PairwiseWithoutSmallAntsFeeder60m = emmeans(Exp460mWithoutSmall_m, spec= "feedertype")

PairwiseWithoutSmallAntsFeeder60m=contrast(PairwiseWithoutSmallAntsFeeder60m, method = "pairwise")
summary(PairwiseWithoutSmallAntsFeeder60m)
##  contrast     estimate    SE  df t.ratio p.value
##  dead - novel   -0.403 0.052 266  -7.751  <.0001
##
## Results are averaged over the levels of: Dead_odour
## Results are given on the log (not the response) scale.
#Glmm Ants staying/feeding on the two different feeders over 23h

Exp423hWithoutSmall_m <-glmmTMB(antsOnFeeder ~ feedertype + Time_after_start_min + Dead_odour + (1|Colony), ziformula = ~1,
                              family= poisson,
                              data= Exp423hWithoutSmall)
## Warning in (function (start, objective, gradient = NULL, hessian = NULL, : NA/
## NaN function evaluation
summary(Exp423hWithoutSmall_m)
##  Family: poisson  ( log )
## Formula:
## antsOnFeeder ~ feedertype + Time_after_start_min + Dead_odour +
##     (1 | Colony)
## Zero inflation:                ~1
## Data: Exp423hWithoutSmall
##
##      AIC      BIC   logLik deviance df.resid
##    474.7    498.7   -231.3    462.7      396
##
## Random effects:
##
## Conditional model:
##  Groups Name        Variance Std.Dev.
##  Colony (Intercept) 0.4977   0.7055
## Number of obs: 402, groups:  Colony, 8
##
## Conditional model:
##                        Estimate Std. Error z value Pr(>|z|)
## (Intercept)          -0.2377081  0.3860030  -0.616 0.538014
## feedertypenovel       0.2706608  0.2327537   1.163 0.244885
## Time_after_start_min -0.0016647  0.0002926  -5.689 1.28e-08 ***
## Dead_odourStrawberry  0.9482599  0.2604722   3.641 0.000272 ***
## ---
## Signif. codes:  0 '***' 0.001 '**' 0.01 '*' 0.05 '.' 0.1 ' ' 1
##
## Zero-inflation model:
##             Estimate Std. Error z value Pr(>|z|)
## (Intercept)   0.1677     0.2850   0.588    0.556
dharmitExp423hWithoutSmall <-simulateResiduals(Exp423hWithoutSmall_m)
plot(dharmitExp423hWithoutSmall)
```

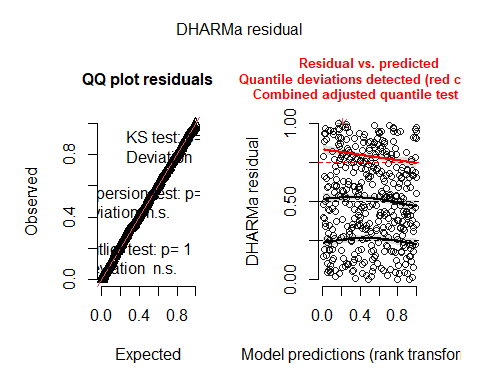

```
# Pairwise
PairwiseWithoutSmallAntsFeeder24h = emmeans(Exp423hWithoutSmall_m, spec= "feedertype")

PairwiseWithoutSmallAntsFeeder24h =contrast(PairwiseWithoutSmallAntsFeeder24h , method = "pairwise")
summary(PairwiseWithoutSmallAntsFeeder24h )
##  contrast     estimate    SE  df t.ratio p.value
##  dead - novel   -0.271 0.233 396  -1.163  0.2456
##
## Results are averaged over the levels of: Dead_odour
## Results are given on the log (not the response) scale.
# GLMM Proportion of ants feeding over 60m on a specific feeder

# Because there is later a plot issue we need to do a subdata  again

Exp4WithoutSmallM <- read_excel("C:/Users/LocalAdmin/Documents/ThomasWagnerPhD/PhD/RawData/OvercomingNegativStimuli/All4Exp/Exp4Collective/Exp4Recovery.xlsx")


Exp4WithoutSmallM$feedertype <- as.factor(Exp4WithoutSmallM$feedertype)
Exp4WithoutSmallM$Colony <- as.factor(Exp4WithoutSmallM$Colony)

Exp4WithoutSmallM$antsOnFeeder <- as.numeric(Exp4WithoutSmallM$antsOnFeeder)
Exp4WithoutSmallM$prop.on.feeder <- as.numeric(Exp4WithoutSmallM$prop.on.feeder)
## Warning: NAs durch Umwandlung erzeugt
Exp4WithoutSmallM <- subset(Exp4WithoutSmallM, !is.infinite(prop.on.feeder))

Exp4WithoutSmallM <- na.omit(Exp4WithoutSmallM)

Exp4WithoutSmallM$Colony <- as.numeric(gsub("[^0-9]", "", Exp4WithoutSmallM$Colony))


Exp4WithoutSmall60mM<-subset(Exp4WithoutSmallM, Time_after_start_min < 120 & total.feeding.ants.whole.exp > 40)

Exp423hWithoutSmall23hM <-subset(Exp4WithoutSmallM , Time_after_start_min > 119 & total.feeding.ants.whole.exp > 40)


Exp4WithoutSmall60mM_m <- gamlss(prop.on.feeder ~ feedertype + Time_after_start_min + Dead_odour,
                                             family = BEINF,
                                             data = Exp4WithoutSmall60mM)
## GAMLSS-RS iteration 1: Global Deviance = 508.7339
## GAMLSS-RS iteration 2: Global Deviance = 508.5392
## GAMLSS-RS iteration 3: Global Deviance = 508.5387
summary(Exp4WithoutSmall60mM_m)
## ******************************************************************
## Family:  c("BEINF", "Beta Inflated")
##
## Call:  gamlss(formula = prop.on.feeder ~ feedertype + Time_after_start_min +
##     Dead_odour, family = BEINF, data = Exp4WithoutSmall60mM)
##
## Fitting method: RS()
##
## ------------------------------------------------------------------
## Mu link function:  logit
## Mu Coefficients:
##                        Estimate Std. Error t value Pr(>|t|)
## (Intercept)          -2.513e-01  2.250e-01  -1.117  0.26502
## feedertypenovel       5.027e-01  1.729e-01   2.907  0.00396 **
## Time_after_start_min  1.238e-18  5.068e-03   0.000  1.00000
## Dead_odourStrawberry  2.722e-16  1.739e-01   0.000  1.00000
## ---
## Signif. codes:  0 '***' 0.001 '**' 0.01 '*' 0.05 '.' 0.1 ' ' 1
##
## ------------------------------------------------------------------
## Sigma link function:  logit
## Sigma Coefficients:
##             Estimate Std. Error t value Pr(>|t|)
## (Intercept)  0.21301    0.07567   2.815  0.00525 **
## ---
## Signif. codes:  0 '***' 0.001 '**' 0.01 '*' 0.05 '.' 0.1 ' ' 1
##
## ------------------------------------------------------------------
## Nu link function:  log
## Nu Coefficients:
##             Estimate Std. Error t value Pr(>|t|)
## (Intercept)  -1.0118     0.1561  -6.484  4.5e-10 ***
## ---
## Signif. codes:  0 '***' 0.001 '**' 0.01 '*' 0.05 '.' 0.1 ' ' 1
##
## ------------------------------------------------------------------
## Tau link function:  log
## Tau Coefficients:
##             Estimate Std. Error t value Pr(>|t|)
## (Intercept)   -1.012      0.156  -6.483 4.52e-10 ***
## ---
## Signif. codes:  0 '***' 0.001 '**' 0.01 '*' 0.05 '.' 0.1 ' ' 1
##
## ------------------------------------------------------------------
## No. of observations in the fit:  266
## Degrees of Freedom for the fit:  7
##       Residual Deg. of Freedom:  259
##                       at cycle:  3
##
## Global Deviance:     508.5387
##             AIC:     522.5387
##             SBC:     547.6232
## ******************************************************************
# % of ants sticking on a feeder

proportions <- with(Exp4WithoutSmall60mM, prop.table(table(feedertype)))

# Print the proportions
print(proportions)
## feedertype
##  dead novel
##   0.5   0.5
# Pairwise
PairwiseWithoutSmall60m = emmeans(Exp4WithoutSmall60mM_m, spec= "feedertype")
## ******************************************************************
## Family:  c("BEINF", "Beta Inflated")
##
## Call:  gamlss(formula = prop.on.feeder ~ feedertype + Time_after_start_min +
##     Dead_odour, family = BEINF, data = Exp4WithoutSmall60mM)
##
## Fitting method: RS()
##
## ------------------------------------------------------------------
## Mu link function:  logit
## Mu Coefficients:
##                        Estimate Std. Error t value Pr(>|t|)
## (Intercept)          -2.513e-01  2.250e-01  -1.117  0.26502
## feedertypenovel       5.027e-01  1.729e-01   2.907  0.00396 **
## Time_after_start_min  1.238e-18  5.068e-03   0.000  1.00000
## Dead_odourStrawberry  2.722e-16  1.739e-01   0.000  1.00000
## ---
## Signif. codes:  0 '***' 0.001 '**' 0.01 '*' 0.05 '.' 0.1 ' ' 1
##
## ------------------------------------------------------------------
## Sigma link function:  logit
## Sigma Coefficients:
##             Estimate Std. Error t value Pr(>|t|)
## (Intercept)  0.21301    0.07567   2.815  0.00525 **
## ---
## Signif. codes:  0 '***' 0.001 '**' 0.01 '*' 0.05 '.' 0.1 ' ' 1
##
## ------------------------------------------------------------------
## Nu link function:  log
## Nu Coefficients:
##             Estimate Std. Error t value Pr(>|t|)
## (Intercept)  -1.0118     0.1561  -6.484  4.5e-10 ***
## ---
## Signif. codes:  0 '***' 0.001 '**' 0.01 '*' 0.05 '.' 0.1 ' ' 1
##
## ------------------------------------------------------------------
## Tau link function:  log
## Tau Coefficients:
##             Estimate Std. Error t value Pr(>|t|)
## (Intercept)   -1.012      0.156  -6.483 4.52e-10 ***
## ---
## Signif. codes:  0 '***' 0.001 '**' 0.01 '*' 0.05 '.' 0.1 ' ' 1
##
## ------------------------------------------------------------------
## No. of observations in the fit:  266
## Degrees of Freedom for the fit:  7
##       Residual Deg. of Freedom:  259
##                       at cycle:  3
##
## Global Deviance:     508.5387
##             AIC:     522.5387
##             SBC:     547.6232
## ******************************************************************
PairwiseWithoutSmall60m=contrast(PairwiseWithoutSmall60m, method = "pairwise")
summary(PairwiseWithoutSmall60m)
##  contrast     estimate    SE  df z.ratio p.value
##  dead - novel   -0.503 0.173 Inf  -2.907  0.0036
##
## Results are averaged over the levels of: Dead_odour
## Results are given on the log odds ratio (not the response) scale.
# GLMM Proportion of ants feeding over 23h on a specific feeder

Exp4WithoutSmall23hM_m <- gamlss(prop.on.feeder ~ feedertype + Time_after_start_min + Dead_odour,
                                             family = BEINF,
                                             data = Exp423hWithoutSmall23hM)
## GAMLSS-RS iteration 1: Global Deviance = 248.0721
## GAMLSS-RS iteration 2: Global Deviance = 226.8458
## GAMLSS-RS iteration 3: Global Deviance = 222.7846
## GAMLSS-RS iteration 4: Global Deviance = 221.5562
## GAMLSS-RS iteration 5: Global Deviance = 221.1132
## GAMLSS-RS iteration 6: Global Deviance = 220.9385
## GAMLSS-RS iteration 7: Global Deviance = 220.8659
## GAMLSS-RS iteration 8: Global Deviance = 220.8348
## GAMLSS-RS iteration 9: Global Deviance = 220.8212
## GAMLSS-RS iteration 10: Global Deviance = 220.8152
## GAMLSS-RS iteration 11: Global Deviance = 220.8125
## GAMLSS-RS iteration 12: Global Deviance = 220.8113
## GAMLSS-RS iteration 13: Global Deviance = 220.8107
summary(Exp4WithoutSmall23hM_m)
## ******************************************************************
## Family:  c("BEINF", "Beta Inflated")
##
## Call:  gamlss(formula = prop.on.feeder ~ feedertype + Time_after_start_min +
##     Dead_odour, family = BEINF, data = Exp423hWithoutSmall23hM)
##
## Fitting method: RS()
##
## ------------------------------------------------------------------
## Mu link function:  logit
## Mu Coefficients:
##                        Estimate Std. Error t value Pr(>|t|)
## (Intercept)          -2.871e-01  3.635e-01  -0.790    0.431
## feedertypenovel       5.742e-01  3.780e-01   1.519    0.131
## Time_after_start_min -1.060e-18  7.314e-04   0.000    1.000
## Dead_odourStrawberry  3.818e-16  4.117e-01   0.000    1.000
##
## ------------------------------------------------------------------
## Sigma link function:  logit
## Sigma Coefficients:
##             Estimate Std. Error t value Pr(>|t|)
## (Intercept)  -0.7292     0.2563  -2.845  0.00526 **
## ---
## Signif. codes:  0 '***' 0.001 '**' 0.01 '*' 0.05 '.' 0.1 ' ' 1
##
## ------------------------------------------------------------------
## Nu link function:  log
## Nu Coefficients:
##             Estimate Std. Error t value Pr(>|t|)
## (Intercept)   1.5157     0.3179   4.767 5.51e-06 ***
## ---
## Signif. codes:  0 '***' 0.001 '**' 0.01 '*' 0.05 '.' 0.1 ' ' 1
##
## ------------------------------------------------------------------
## Tau link function:  log
## Tau Coefficients:
##             Estimate Std. Error t value Pr(>|t|)
## (Intercept)   1.5169     0.3179   4.772 5.41e-06 ***
## ---
## Signif. codes:  0 '***' 0.001 '**' 0.01 '*' 0.05 '.' 0.1 ' ' 1
##
## ------------------------------------------------------------------
## No. of observations in the fit:  122
## Degrees of Freedom for the fit:  7
##       Residual Deg. of Freedom:  115
##                       at cycle:  13
##
## Global Deviance:     220.8107
##             AIC:     234.8107
##             SBC:     254.4389
## ******************************************************************
#Pairwise


PairwiseWithoutSmall23h = emmeans(Exp4WithoutSmall23hM_m, spec= "feedertype")
## ******************************************************************
## Family:  c("BEINF", "Beta Inflated")
##
## Call:  gamlss(formula = prop.on.feeder ~ feedertype + Time_after_start_min +
##     Dead_odour, family = BEINF, data = Exp423hWithoutSmall23hM)
##
## Fitting method: RS()
##
## ------------------------------------------------------------------
## Mu link function:  logit
## Mu Coefficients:
##                        Estimate Std. Error t value Pr(>|t|)
## (Intercept)          -2.871e-01  3.635e-01  -0.790    0.431
## feedertypenovel       5.742e-01  3.780e-01   1.519    0.131
## Time_after_start_min -1.060e-18  7.314e-04   0.000    1.000
## Dead_odourStrawberry  3.818e-16  4.117e-01   0.000    1.000
##
## ------------------------------------------------------------------
## Sigma link function:  logit
## Sigma Coefficients:
##             Estimate Std. Error t value Pr(>|t|)
## (Intercept)  -0.7292     0.2563  -2.845  0.00526 **
## ---
## Signif. codes:  0 '***' 0.001 '**' 0.01 '*' 0.05 '.' 0.1 ' ' 1
##
## ------------------------------------------------------------------
## Nu link function:  log
## Nu Coefficients:
##             Estimate Std. Error t value Pr(>|t|)
## (Intercept)   1.5157     0.3179   4.767 5.51e-06 ***
## ---
## Signif. codes:  0 '***' 0.001 '**' 0.01 '*' 0.05 '.' 0.1 ' ' 1
##
## ------------------------------------------------------------------
## Tau link function:  log
## Tau Coefficients:
##             Estimate Std. Error t value Pr(>|t|)
## (Intercept)   1.5169     0.3179   4.772 5.41e-06 ***
## ---
## Signif. codes:  0 '***' 0.001 '**' 0.01 '*' 0.05 '.' 0.1 ' ' 1
##
## ------------------------------------------------------------------
## No. of observations in the fit:  122
## Degrees of Freedom for the fit:  7
##       Residual Deg. of Freedom:  115
##                       at cycle:  13
##
## Global Deviance:     220.8107
##             AIC:     234.8107
##             SBC:     254.4389
## ******************************************************************
PairwiseWithoutSmall23h=contrast(PairwiseWithoutSmall23h, method = "pairwise")
summary(PairwiseWithoutSmall23h)
##  contrast     estimate    SE  df z.ratio p.value
##  dead - novel   -0.574 0.378 Inf  -1.519  0.1288
##
## Results are averaged over the levels of: Dead_odour
## Results are given on the log odds ratio (not the response) scale.
```

##### Figures for supplement - Plots with all colonies

```
posn.d <- position_dodge(width = 2.5)

Exp4RecoveryOnly60m_summary <- Exp4RecoveryOnly60m %>%
  group_by(Time_after_start_min, feedertype) %>%
  summarise(mean_antsOnFeeder = mean(antsOnFeeder), SE = sd(antsOnFeeder) / sqrt(n()))
## `summarise()` has grouped output by 'Time_after_start_min'. You can override
## using the `.groups` argument.
fig4Curve60mWithAllColonies <- ggplot(data = Exp4RecoveryOnly60m_summary, mapping = aes(x = Time_after_start_min, y = mean_antsOnFeeder, fill = feedertype)) +
  geom_point(aes(color = feedertype), shape = 16, size = 3, stroke = 0.8) +
  geom_line(aes(color = feedertype), size = 1) +
  scale_y_continuous(limits = c(0, NA)) +
  labs(x = "Time after observation start (minutes)", y = "Ants on feeder") +
  scale_fill_manual(name = "Feeder type", values = c("red", "dodgerblue"), labels = c("corpse", "novel")) +
  scale_color_manual(name = "Feeder type", values = c("red", "dodgerblue"), labels = c("corpse", "novel")) +
  theme_bw() +
  scale_x_continuous(breaks = seq(0, 60, by = 5)) +
  theme(
    axis.line = element_line(color = "black"),
    plot.background = element_blank(),
    panel.grid.major = element_blank(),
    panel.grid.minor = element_blank(),
    panel.border = element_blank(),
    axis.text = element_text(size = 14),
    axis.title = element_text(size = 15, face = "bold"),
    legend.position = "none"  # Remove the legend
  )

print(fig4Curve60mWithAllColonies)
```

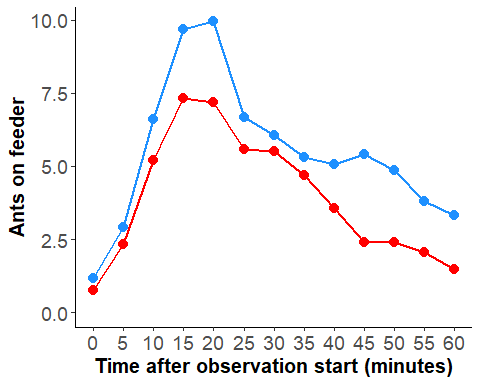

```
ggsave("fig4Curve60mWithAllColonies.png", plot = fig4Curve60mWithAllColonies, dpi = 300, width = 8, height = 6)


# Curve plot "Number of ants on different feeder types for following 23h with all colonies" #


posn.d2 <- position_dodge(width = 0.5)

Exp4Recovery23hours_summary <- Exp4Recovery23h %>%
  group_by(Time_after_start_min, feedertype) %>%
  summarise(mean_antsOnFeeder = mean(antsOnFeeder), .groups = "keep")

fig4Curve23hWithAllColonies  <- ggplot(data = Exp4Recovery23hours_summary, mapping = aes(x = Time_after_start_min / 60, y = mean_antsOnFeeder, color = feedertype, group = feedertype)) +
  geom_point(position = posn.d2, shape = 16, size = 3, stroke = 0.8) +
  geom_line(size = 1) +
  labs(x = "Time after observation start (hours)", y = "Ants on feeder") +
  scale_x_continuous(breaks = seq(2, 24, by = 1), labels = seq(2, 24, by = 1)) +
  scale_color_manual(name = "Feeder Type", values = c("red", "dodgerblue"), labels = c("corpse", "novel")) +
  theme_bw() +
  theme(
    axis.line = element_line(color = "black"),
    plot.background = element_blank(),
    panel.grid.major = element_blank(),
    panel.grid.minor = element_blank(),
    panel.border = element_blank(),
    axis.text = element_text(size = 12),  # Adjust the font size here
    axis.title = element_text(size = 15, face = "bold"),
    legend.title = element_text(size = 14, face = "bold"),
    legend.text = element_text(size = 12),
    legend.position = "none"  # Remove the legend
  )

print(fig4Curve23hWithAllColonies)
```

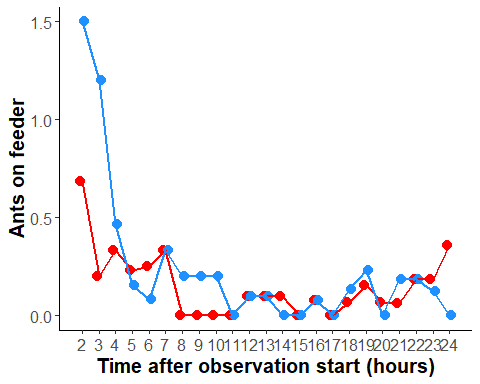

```
ggsave("fig4Curve23hWithAllColonies.png", plot = fig4Curve23hWithAllColonies, dpi = 300, width = 8, height = 6)


#  Bar plot "Number of ants on different feeder types for 60m with all colonies" #


Exp4RecoveryOnly60m_summary$feedertype <- forcats::fct_recode(Exp4RecoveryOnly60m_summary$feedertype, "corpse" = "dead")

fig4Bar60mWithAllColonies <- ggplot(data = Exp4RecoveryOnly60m_summary, aes(x = feedertype, y = mean_antsOnFeeder, fill = feedertype)) +
  stat_summary(fun = "mean", geom = "bar", width = 0.75) +
  stat_summary(fun.data = "mean_cl_boot", geom = "errorbar", width = 0.5) +
  ylab("Ants on feeder") +
  xlab("") +  # Change the x-axis label
  scale_fill_manual(name = "", values = c("red", "dodgerblue"), labels = c("corpse", "novel")) +  # Change the legend name and colors
  coord_cartesian(ylim = c(0, 10)) +  # Set the y-axis limits to 0 and 7.5
  scale_y_continuous(expand = expansion(mult = c(0, 0.1))) +
  theme_classic() +  # Use classic theme
  theme(
    axis.line = element_line(color = "black"),
    plot.background = element_blank(),
    panel.grid.major = element_blank(),
    panel.grid.minor = element_blank(),
    panel.border = element_blank(),
    axis.text = element_text(size = 12),
    axis.title = element_text(size = 15, face = "bold"),legend.position = "none",
    axis.text.x = element_text()
  )

print(fig4Bar60mWithAllColonies)
```

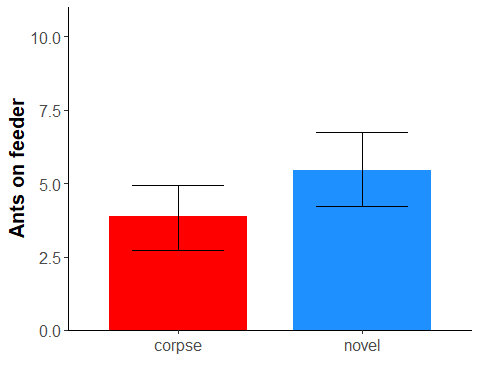

```
ggsave("fig4Bar60mWithAllColonies.png", plot = fig4Bar60mWithAllColonies, dpi = 300, width = 8, height = 6)

#  Bar plot "Number of ants on different feeder types for 23h with all colonies

Exp4Recovery23hours_summary$feedertype <- forcats::fct_recode(Exp4Recovery23hours_summary$feedertype, "corpse" = "dead")

fig4Bar23hWithAllColonies <- ggplot(data = Exp4Recovery23hours_summary, aes(x = feedertype, y = mean_antsOnFeeder, fill = feedertype)) +
  stat_summary(fun = "mean", geom = "bar", width = 0.75) +
  stat_summary(fun.data = "mean_cl_boot", geom = "errorbar", width = 0.5) +
  ylab("Ants on feeder") +
  xlab("") +  # Change the x-axis label
  scale_fill_manual(name = "", values = c("red", "dodgerblue"), labels = c("corpse", "novel")) +  # Change the legend name and colors
  coord_cartesian(ylim = c(0, 1.4)) +  # Set the y-axis limits to 0 and 7.5
  scale_y_continuous(expand = expansion(mult = c(0, 0.1))) +
  theme_classic() +  # Use classic theme
  theme(
    axis.line = element_line(color = "black"),
    plot.background = element_blank(),
    panel.grid.major = element_blank(),
    panel.grid.minor = element_blank(),
    panel.border = element_blank(),
    axis.text = element_text(size = 12),
    axis.title = element_text(size = 15, face = "bold"),legend.position = "none",
    axis.text.x = element_text()
  )

print(fig4Bar23hWithAllColonies)
```

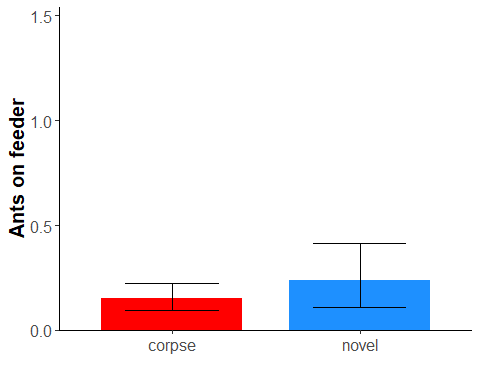

```
ggsave("fig4Bar23hWithAllColonies.png", plot = fig4Bar23hWithAllColonies, dpi = 300, width = 8, height = 6)


# Proportion of ants sticking on one of the feeder over 60m

Exp4Prob60m$feedertype <- forcats::fct_recode(Exp4Prob60m$feedertype, "corpse" = "dead")

Exp4Prob60m_summary <- Exp4Prob60m  %>%
  group_by(Time_after_start_min, feedertype) %>%
  summarise(mean_proportion = mean(prop.on.feeder), SE = sd(prop.on.feeder) / sqrt(n()))
## `summarise()` has grouped output by 'Time_after_start_min'. You can override
## using the `.groups` argument.
fig4Bar60mWithAllColoniesProp <- ggplot(data = Exp4Prob60m_summary, aes(x = feedertype, y = mean_proportion, fill = feedertype)) +
  stat_summary(fun = "mean", geom = "bar", width = 0.75) +
  stat_summary(fun.data = "mean_cl_boot", geom = "errorbar", width = 0.5) +
  ylab("Proportion of ants on feeder") +
  xlab("") +  # Change the x-axis label
  scale_fill_manual(name = "Feeder Type", values = c("red", "dodgerblue"), labels = c("corpse", "novel")) +  # Change the legend name and colors
  coord_cartesian(ylim = c(0, 1)) +  # Set the y-axis limits to 0 and 1
  scale_y_continuous(expand = expansion(mult = c(0, 0.1))) +
  theme_classic() +  # Use classic theme
  theme(
    axis.line = element_line(color = "black"),
    plot.background = element_blank(),
    panel.grid.major = element_blank(),
    panel.grid.minor = element_blank(),
    panel.border = element_blank(),
    axis.text = element_text(size = 12),
    axis.title = element_text(size = 15, face = "bold"),
    axis.text.x = element_text(),legend.position = "none"
  )

print(fig4Bar60mWithAllColoniesProp)
```

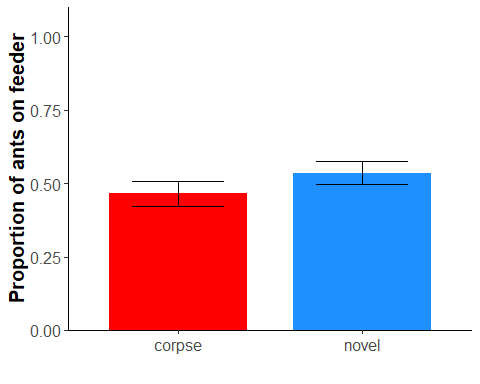

```
ggsave("fig4Bar60mWithAllColoniesProp.png", plot = fig4Bar60mWithAllColoniesProp, dpi = 300, width = 8, height = 6)


# Proportion of ants sticking on one of the feeder over 23h


Exp4Prob23h$feedertype <- forcats::fct_recode(Exp4Prob23h$feedertype, "corpse" = "dead")

Exp4Prob23h_summary <- Exp4Prob23h  %>%
  group_by(Time_after_start_min, feedertype) %>%
  summarise(mean_proportion = mean(prop.on.feeder), SE = sd(prop.on.feeder) / sqrt(n()))
## `summarise()` has grouped output by 'Time_after_start_min'. You can override
## using the `.groups` argument.
fig4Bar23hWithAllColoniesProp <- ggplot(data = Exp4Prob23h_summary, aes(x = feedertype, y = mean_proportion, fill = feedertype)) +
  stat_summary(fun = "mean", geom = "bar", width = 0.75) +
  stat_summary(fun.data = "mean_cl_boot", geom = "errorbar", width = 0.5) +
  ylab("Proportion of ants on feeder") +
  xlab("") +  # Change the x-axis label
  scale_fill_manual(name = "Feeder Type", values = c("red", "dodgerblue"), labels = c("corpse", "novel")) +  # Change the legend name and colors
  coord_cartesian(ylim = c(0, 1)) +  # Set the y-axis limits to 0 and 1
  scale_y_continuous(expand = expansion(mult = c(0, 0.1))) +
  theme_classic() +  # Use classic theme
  theme(
    axis.line = element_line(color = "black"),
    plot.background = element_blank(),
    panel.grid.major = element_blank(),
    panel.grid.minor = element_blank(),
    panel.border = element_blank(),
    axis.text = element_text(size = 12),
    axis.title = element_text(size = 15, face = "bold"),
    axis.text.x = element_text(),legend.position = "none"
  )

print(fig4Bar23hWithAllColoniesProp)
```

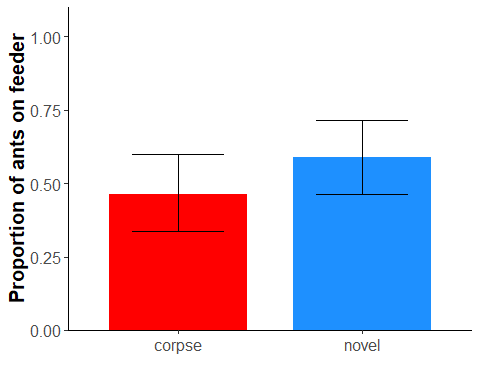

ggsave("fig4Bar23hWithAllColoniesProp.png", plot = fig4Bar23hWithAllColoniesProp, dpi = 300, width = 8, height = 6)

##### Figures for the manuscript - Plots without the very small colonies.

```
# Curve plot "Number of ants on different feeder types within 60m

posn.d <- position_dodge(width = 2.5)

Exp4Only60m_summary_Without <- Exp460mWithoutSmall %>%
  group_by(Time_after_start_min, feedertype) %>%
  summarise(mean_antsOnFeeder = mean(antsOnFeeder), SE = sd(antsOnFeeder) / sqrt(n()))
## `summarise()` has grouped output by 'Time_after_start_min'. You can override
## using the `.groups` argument.
fig4Curve60mWithout <- ggplot(data = Exp4Only60m_summary_Without, mapping = aes(x = Time_after_start_min, y = mean_antsOnFeeder, fill = feedertype)) +
  geom_point(aes(color = feedertype), shape = 16, size = 3, stroke = 0.8) +
  geom_line(aes(color = feedertype), size = 1) +
  coord_cartesian(ylim = c(0, 16)) +
  scale_y_continuous(limits = c(0, 16)) +
  labs(x = "Time after observation start (minutes)", y = "Ants on feeder") +
  scale_fill_manual(name = "Feeder type", values = c("red", "dodgerblue"), labels = c("corpse", "novel")) +
  scale_color_manual(name = "Feeder type", values = c("red", "dodgerblue"), labels = c("corpse", "novel")) +
  theme_bw() +
  scale_x_continuous(breaks = seq(0, 60, by = 5)) +
  theme(
    axis.line = element_line(color = "black"),
    plot.background = element_blank(),
    panel.grid.major = element_blank(),
    panel.grid.minor = element_blank(),
    panel.border = element_blank(),
    axis.text = element_text(size = 16),
    axis.title = element_text(size = 20, face = "bold"),
    legend.position = "none"  # Remove the legend
  )

print(fig4Curve60mWithout)
```

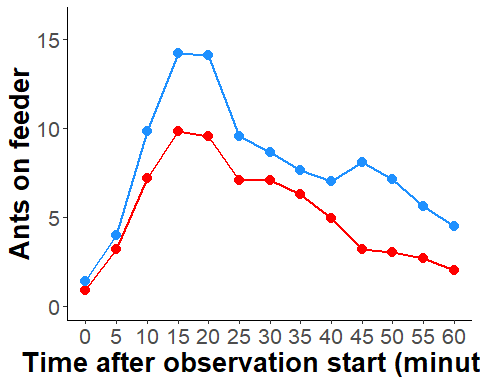

```
ggsave("fig4Curve60mWithout.png", plot = fig4Curve60mWithout, dpi = 300, width = 8, height = 6)

# Curve plot "Number of ants on different feeder types for following 23h


posn.d2 <- position_dodge(width = 0.5)

Exp4Only23h_summary_Without <- Exp423hWithoutSmall %>%
  group_by(Time_after_start_min, feedertype) %>%
  summarise(mean_antsOnFeeder = mean(antsOnFeeder), .groups = "keep")

fig4Curve623hWithout  <- ggplot(data = Exp4Only23h_summary_Without, mapping = aes(x = Time_after_start_min / 60, y = mean_antsOnFeeder, color = feedertype, group = feedertype)) +
  geom_point(position = posn.d2, shape = 16, size = 3, stroke = 0.8) +
  geom_line(size = 1) +
  labs(x = "Time after observation start (hours)", y = "Ants on feeder") +
   coord_cartesian(ylim = c(0, 2)) +
  scale_x_continuous(breaks = seq(2, 24, by = 1), labels = seq(2, 24, by = 1)) +
  scale_color_manual(name = "Feeder Type", values = c("red", "dodgerblue"), labels = c("corpse", "novel")) +
  theme_bw() +
  theme(
    axis.line = element_line(color = "black"),
    plot.background = element_blank(),
    panel.grid.major = element_blank(),
    panel.grid.minor = element_blank(),
    panel.border = element_blank(),
    axis.text = element_text(size = 16),  # Adjust the font size here
    axis.title = element_text(size = 20, face = "bold"),
    legend.title = element_text(size = 20, face = "bold"),
    legend.text = element_text(size = 18),
    legend.position = "none"  # Remove the legend
  )

print(fig4Curve623hWithout)
```

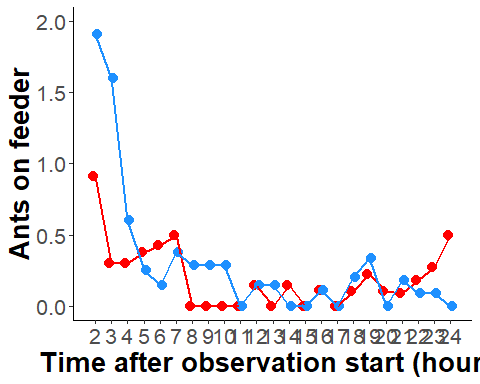

```
ggsave("Fig4Curve623hWithout.png", plot = fig4Curve23hWithAllColonies, dpi = 300, width = 8, height = 6)


#  Bar plot "Number of ants on different feeder types for 60m without small colonies.


Exp460mWithoutSmall$feedertype <- forcats::fct_recode(Exp460mWithoutSmall$feedertype, "corpse" = "dead")


fig4Bar60mWithout <- ggplot(data = Exp460mWithoutSmall, aes(x = feedertype, y = antsOnFeeder, fill = feedertype)) +
  stat_summary(fun = "mean", geom = "bar", width = 0.75) +
  stat_summary(fun.data = "mean_cl_boot", geom = "errorbar", width = 0.5) +
   ylab("Ants on feeder") +
  xlab("") +  # # Change the x-axis label
  scale_fill_manual(name = "Feeder Type", values = c("red", "dodgerblue"), labels = c("corpse", "novel")) +  # Change the legend name and colors
  coord_cartesian(ylim = c(0, 16)) +  # Set the y-axis limits to 0 and 7.5
  scale_y_continuous(expand = expansion(mult = c(0, 0.1))) +
  theme_classic() +  # Use classic theme
  theme(
    axis.line = element_line(color = "black"),
    plot.background = element_blank(),
    panel.grid.major = element_blank(),
    panel.grid.minor = element_blank(),
    panel.border = element_blank(),
    axis.text = element_text(size = 16),
    axis.title = element_text(size = 20, face = "bold"),
    axis.text.x = element_text(16),
    legend.text = element_text(size = 18),    # Adjust the font size of the legend text
    legend.title = element_text(size = 18, face = "bold")

  )

# Print the plot with the legend
print(fig4Bar60mWithout)
## Warning in grid.Call(C_stringMetric, as.graphicsAnnot(x$label)):
## Zeichensatzfamilie in der Windows Zeichensatzdatenbank nicht gefunden
## Warning in grid.Call(C_textBounds, as.graphicsAnnot(x$label), x$x, x$y, :
## Zeichensatzfamilie in der Windows Zeichensatzdatenbank nicht gefunden

## Warning in grid.Call(C_textBounds, as.graphicsAnnot(x$label), x$x, x$y, :
## Zeichensatzfamilie in der Windows Zeichensatzdatenbank nicht gefunden
```

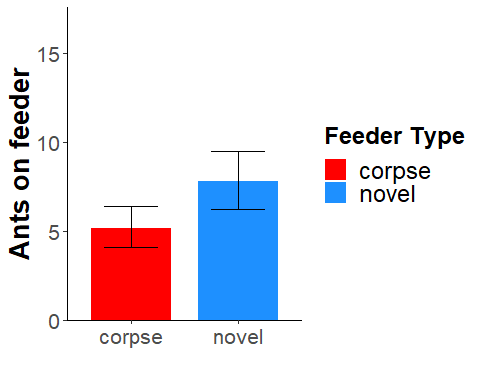

```
print(fig4Bar60mWithout)
## Warning in grid.Call(C_textBounds, as.graphicsAnnot(x$label), x$x, x$y, :
## Zeichensatzfamilie in der Windows Zeichensatzdatenbank nicht gefunden

## Warning in grid.Call(C_textBounds, as.graphicsAnnot(x$label), x$x, x$y, :
## Zeichensatzfamilie in der Windows Zeichensatzdatenbank nicht gefunden
```

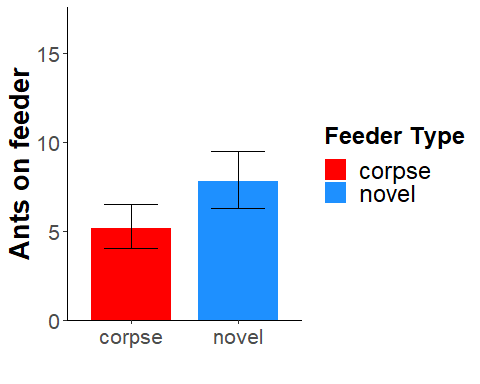

```
ggsave("fig4Bar60mWithout.png", plot = fig4Bar60mWithout, dpi = 300, width = 8, height = 6)
## Warning in grid.Call(C_textBounds, as.graphicsAnnot(x$label), x$x, x$y, :
## Zeichensatzfamilie in der Windows Zeichensatzdatenbank nicht gefunden

## Warning in grid.Call(C_textBounds, as.graphicsAnnot(x$label), x$x, x$y, :
## Zeichensatzfamilie in der Windows Zeichensatzdatenbank nicht gefunden
#  Bar plot "Number of ants on different feeder types for the following 23h without small colonies.

Exp423hWithoutSmall$feedertype <- forcats::fct_recode(Exp423hWithoutSmall$feedertype, "corpse" = "dead")

fig4Bar23hWithout <- ggplot(data = Exp423hWithoutSmall, aes(x = feedertype, y = antsOnFeeder, fill = feedertype)) +
  stat_summary(fun = "mean", geom = "bar", width = 0.75) +
  stat_summary(fun.data = "mean_cl_boot", geom = "errorbar", width = 0.5) +
  ylab("Ants on feeder") +
  xlab("") +  # Set x-axis title to be empty
  scale_fill_manual(name = "Feeder Type", values = c("red", "dodgerblue"), labels = c("corpse", "novel")) +  # Change the legend name and colors
  coord_cartesian(ylim = c(0, 2)) +  # Set the y-axis limits to 0 and 2
  scale_y_continuous(expand = expansion(mult = c(0, 0.1))) +
  theme_classic() +  # Use classic theme
  theme(
    axis.line = element_line(color = "black"),
    plot.background = element_blank(),
    panel.grid.major = element_blank(),
    panel.grid.minor = element_blank(),
    panel.border = element_blank(),
    axis.text = element_text(size = 16),      # Adjust the font size for both x and y axes
    axis.title = element_text(size = 20, face = "bold"),
    axis.text.x = element_text(size = 16),    # Adjust the font size for x-axis labels
    legend.text = element_text(size = 18),    # Adjust the font size of the legend text
    legend.title = element_text(size = 18, face = "bold")  # Adjust the font size and font weight for the legend title
  )

# Print the plot with the legend
print(fig4Bar23hWithout)
```

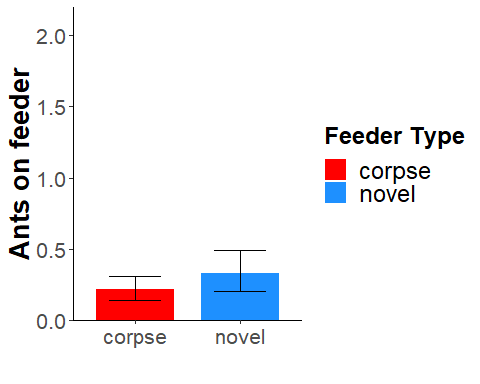

```
ggsave("fig4Bar23hWithout.png", plot = fig4Bar23hWithout, dpi = 300, width = 8, height = 6)


# Proportion of ants sticking on one of the feeder over 60m without small colonies#

Exp4WithoutSmall60mM$feedertype <- forcats::fct_recode(Exp4WithoutSmall60mM$feedertype, "corpse" = "dead")

Exp4Prob60mWithout_summary <- Exp4WithoutSmall60mM  %>%
  group_by(Time_after_start_min, feedertype) %>%
  summarise(mean_proportion = mean(prop.on.feeder), SE = sd(prop.on.feeder) / sqrt(n()))
## `summarise()` has grouped output by 'Time_after_start_min'. You can override
## using the `.groups` argument.
fig4Bar60mWithoutProp <- ggplot(data = Exp4Prob60mWithout_summary, aes(x = feedertype, y = mean_proportion, fill = feedertype)) +
  stat_summary(fun = "mean", geom = "bar", width = 0.75) +
  stat_summary(fun.data = "mean_cl_boot", geom = "errorbar", width = 0.5) +
  ylab("Proportion of ants on feeder") +
  xlab("") +  # Change the x-axis label
  scale_fill_manual(name = "Feeder Type", values = c("red", "dodgerblue"), labels = c("corpse", "novel")) +  # Change the legend name and colors
  coord_cartesian(ylim = c(0, 1)) +  # Set the y-axis limits to 0 and 1
  scale_y_continuous(expand = expansion(mult = c(0, 0.1))) +
  theme_classic() +  # Use classic theme
  theme(
    axis.line = element_line(color = "black"),
    plot.background = element_blank(),
    panel.grid.major = element_blank(),
    panel.grid.minor = element_blank(),
    panel.border = element_blank(),
    axis.text = element_text(size = 12),
    axis.title = element_text(size = 20, face = "bold"),
    axis.text.x = element_text(),legend.position = "none"
  )

print(fig4Bar60mWithoutProp)
```

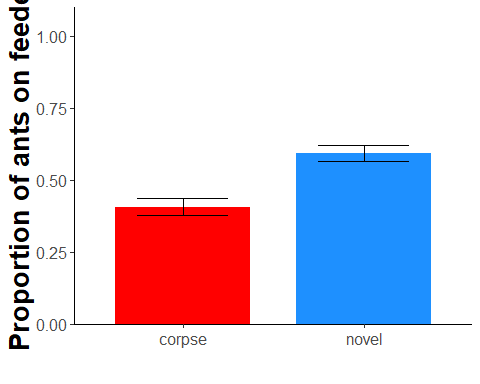

```
ggsave("fig4Bar60mWithoutProp.png", plot = fig4Bar60mWithoutProp, dpi = 300, width = 8, height = 6)


# Proportion of ants sticking on one of the feeder over the following 23h without small colonies

Exp423hWithoutSmall23hM$feedertype <- forcats::fct_recode(Exp423hWithoutSmall23hM$feedertype, "corpse" = "dead")
Exp4Prob23hWithout_summary <- Exp423hWithoutSmall23hM %>%
  group_by(Time_after_start_min, feedertype) %>%
  summarise(mean_proportion = mean(prop.on.feeder), SE = sd(prop.on.feeder) / sqrt(n()))
## `summarise()` has grouped output by 'Time_after_start_min'. You can override
## using the `.groups` argument.
fig4Bar23hWithoutProp <- ggplot(data = Exp4Prob23hWithout_summary, aes(x = feedertype, y = mean_proportion, fill = feedertype)) +
  stat_summary(fun = "mean", geom = "bar", width = 0.75) +
  stat_summary(fun.data = "mean_cl_boot", geom = "errorbar", width = 0.5) +
  ylab("Proportion of ants on feeder") +
  xlab("") +  # Change the x-axis label
  scale_fill_manual(name = "Feeder Type", values = c("red", "dodgerblue"), labels = c("corpse", "novel")) +  # Change the legend name and colors
  coord_cartesian(ylim = c(0, 1)) +  # Set the y-axis limits to 0 and 1
  scale_y_continuous(expand = expansion(mult = c(0, 0.1))) +
  theme_classic() +  # Use classic theme
  theme(
    axis.line = element_line(color = "black"),
    plot.background = element_blank(),
    panel.grid.major = element_blank(),
    panel.grid.minor = element_blank(),
    panel.border = element_blank(),
    axis.text = element_text(size = 12),
    axis.title = element_text(size = 20, face = "bold"),
    axis.text.x = element_text(),legend.position = "none"

  )


print(fig4Bar23hWithoutProp)
```

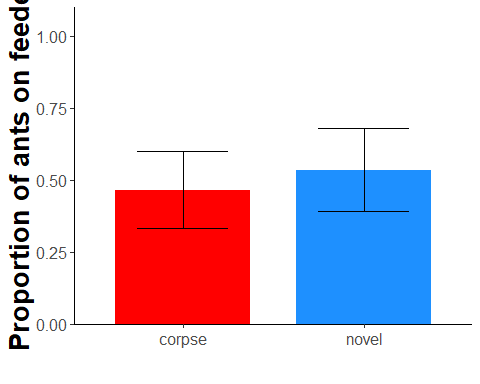

ggsave("fig4Bar23hWithoutProp.png", plot = fig4Bar23hWithoutProp, dpi = 300, width = 8, height = 6)

#Merging all 6 Panels into 1 figure - With all colonies for supplement

```
# Define the desired heights for each row
curve_height <- 2.5
bar_height <- 0.8

# Combine curve plots and bar plots separately
row_1 <- grid.arrange(
  fig4Curve60mWithAllColonies, fig4Bar60mWithAllColonies, fig4Bar60mWithAllColoniesProp,
  ncol = 3, widths = c(curve_height, bar_height, bar_height)
)
```

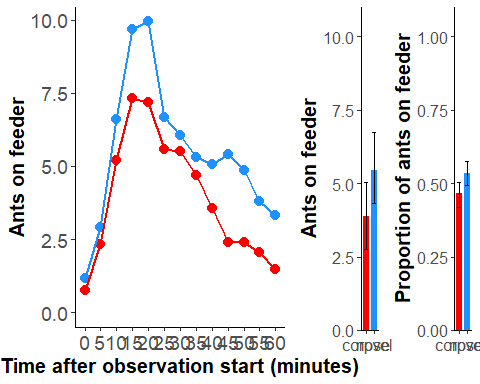

```
row_2 <- grid.arrange(
  fig4Curve23hWithAllColonies, fig4Bar23hWithAllColonies, fig4Bar23hWithAllColoniesProp,
  ncol = 3, widths = c(curve_height, bar_height, bar_height)
)
```

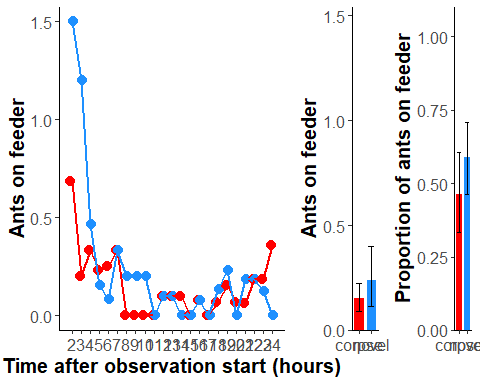

```
# Combine the rows into a 2x3 grid
MergedExp4WithAllColoniesProp <- grid.arrange(
  row_1, row_2,
  nrow = 2
)
```

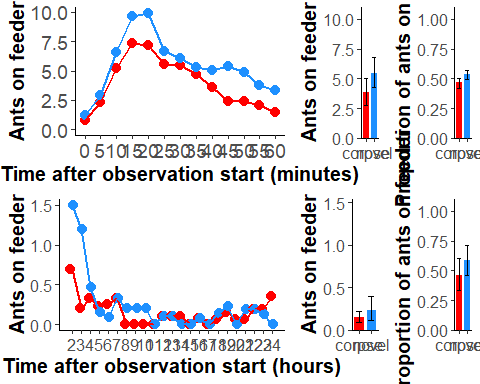

```
ggsave("MergedExp4WithAllColoniesProp.png", plot = MergedExp4WithAllColoniesProp, dpi = 300, width = 12, height = 7)


ppt2 <- read_pptx()

# Add a slide with a title
slide_layout <- "Title and Content"  # You can change the slide layout if needed
ppt2 <- add_slide(ppt2, layout = slide_layout, master = "Office Theme")

# Set the title for the slide
title <- "MergedExp4WithAllColoniesProp"  # Change the title as needed
ppt2 <- ph_with(ppt2, value = title, location = ph_location(type = "title", index = 1))

# Save the combined_fig as a temporary image
temp_img_file <- tempfile(fileext = ".png")
ggsave(temp_img_file, MergedExp4WithAllColoniesProp, width = 12, height = 7)  # Adjust width and height if needed

# Insert the image into the slide
ppt2 <- ph_with(ppt2, value = external_img(temp_img_file), location = ph_location_fullsize())

# Save the PowerPoint file
print(ppt2, target = "MergedExp4WithAllColoniesProp.pptx")

# Browse the file
browseURL("MergedExp4WithAllColoniesProp.pptx")
```

#Merging all 6 Panels into 1 figure - without small colonies for manuscript

```
widths <- c(2/3, 1/3)

MergedExp4WithoutSmallColonies <- grid.arrange(
  fig4Curve60mWithout, fig4Bar60mWithout,
  fig4Curve623hWithout, fig4Bar23hWithout,
  ncol = 2,
  widths = widths
)
## Warning in grid.Call(C_textBounds, as.graphicsAnnot(x$label), x$x, x$y, :
## Zeichensatzfamilie in der Windows Zeichensatzdatenbank nicht gefunden

## Warning in grid.Call(C_textBounds, as.graphicsAnnot(x$label), x$x, x$y, :
## Zeichensatzfamilie in der Windows Zeichensatzdatenbank nicht gefunden
```

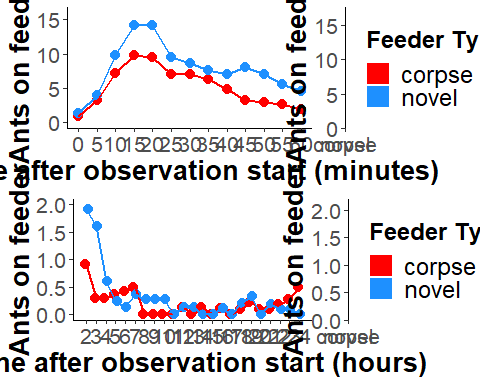

```
ggsave("MergedExp4WithoutSmallColoniesv2.png", plot = MergedExp4WithoutSmallColonies, dpi = 300, width = 16, height = 9)
## Warning in grid.Call(C_textBounds, as.graphicsAnnot(x$label), x$x, x$y, :
## Zeichensatzfamilie in der Windows Zeichensatzdatenbank nicht gefunden

## Warning in grid.Call(C_textBounds, as.graphicsAnnot(x$label), x$x, x$y, :
## Zeichensatzfamilie in der Windows Zeichensatzdatenbank nicht gefunden
# Create a PowerPoint object
ppt <- read_pptx()

# Add a slide with a title
slide_layout <- "Title and Content"  # You can change the slide layout if needed
ppt <- add_slide(ppt, layout = slide_layout, master = "Office Theme")

# Set the title for the slide
title <- "MergedExp4WithoutSmallColoniesv2"  # Change the title as needed
ppt <- ph_with(ppt, value = title, location = ph_location(type = "title", index = 1))

# Save the combined_fig as a temporary image
temp_img_file <- tempfile(fileext = ".png")
ggsave(temp_img_file, MergedExp4WithoutSmallColonies, width = 20, height = 9)  # Adjust width and height if needed
## Warning in grid.Call(C_textBounds, as.graphicsAnnot(x$label), x$x, x$y, :
## Zeichensatzfamilie in der Windows Zeichensatzdatenbank nicht gefunden

## Warning in grid.Call(C_textBounds, as.graphicsAnnot(x$label), x$x, x$y, :
## Zeichensatzfamilie in der Windows Zeichensatzdatenbank nicht gefunden
# Insert the image into the slide
ppt <- ph_with(ppt, value = external_img(temp_img_file), location = ph_location_fullsize())

# Save the PowerPoint file
print(ppt, target = "MergedExp4WithoutSmallColoniesv2.pptx")

# Browse the file
browseURL("MergedExp4WithoutSmallColoniesV2.pptx")


# Proportion only #
Exp4WithoutProp <- grid.arrange(fig4Bar60mWithoutProp, fig4Bar23hWithoutProp, nrow = 1)
```

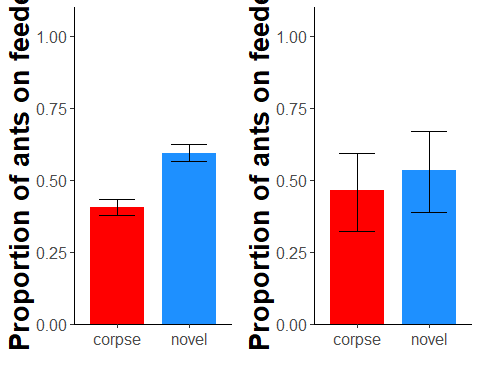

```
print(Exp4WithoutProp)
## TableGrob (1 x 2) "arrange": 2 grobs
##   z     cells    name           grob
## 1 1 (1-1,1-1) arrange gtable[layout]
## 2 2 (1-1,2-2) arrange gtable[layout]
##for powerpoint export##

ggsave ("Exp4WithoutProp.png", plot = Exp4WithoutProp, dpi = 300, width = 20, height = 20, units = c("cm"))

read_pptx() %>%
add_slide(layout = "Title and Content", master = "Office Theme") %>%
  ph_with(value = dml(grid.arrange(fig4Bar60mWithoutProp,fig4Bar23hWithoutProp,nrow = 1)), location = ph_location_type(type = "body")) %>%
 print(target = "Exp4WithoutProp.pptx") %>%
  browseURL()
```
